# An engineered streptavidin condensate platform for chemically inducible control of endogenous proteins in mammalian cells

**DOI:** 10.64898/2026.05.22.725644

**Authors:** Takuya Kamikawa, Cole J. Wilson, Ivy Lan, Yuta Nihongaki

## Abstract

Inducible control of protein activity with temporal precision is essential for understanding and engineering dynamic cellular behaviors. However, current inducible molecular tools largely rely on overexpression of target proteins, which often disrupts the signaling pathways and cellular functions under investigation. A generalizable method to achieve inducible control of endogenous proteins in mammalian cells remains an unmet need. Here, we present a versatile platform based on engineered streptavidin biomolecular condensates to trap and release endogenously tagged proteins. By tagging endogenous loci with a short streptavidin-binding peptide via CRISPR knock-in, our synthetic streptavidin condensates efficiently partition and functionally inhibit the tagged endogenous proteins. The sequestered cargo protein is rapidly released upon the addition of biotin, restoring protein activity within minutes. We demonstrated the broad applicability of this system by controlling diverse endogenous targets: the anterograde motor KIF5B and retrograde motor DYNC1H1, which regulate intracellular vesicle trafficking, and the Arp2/3 complex subunit ARPC3, which regulates actin dynamics. Furthermore, we developed a dual-inducible system based on rapamycin-dependent condensation of streptavidin, enabling both rapid sequestration and release of endogenous proteins at user-defined time points. Altogether, this engineered streptavidin condensate platform provides a robust, rapid, and scalable approach for manipulating endogenous protein function under physiologically relevant conditions in both basic and translational research.

## Introduction

Cells spatially and temporally organize the activities of intracellular biomolecules to execute cellular functions with remarkable precision^1^. While conventional tools such as genetic manipulation and pharmacological perturbation are powerful for identifying key molecular players, they are often insufficient for dissecting the precise spatiotemporal regulation of biomolecules and cellular functions due to their slow kinetics, global effects, and imperfect specificity. Thus, methods enabling rapid manipulation of specific biomolecules at user-defined time points are essential for dissecting spatiotemporal cellular dynamics.

To enable such precise manipulation, researchers have developed inducible molecular tools that regulate target protein activity using controllable domains that undergo chemically or optically induced proximity or allosteric changes^1–3^. While such tools have greatly expanded our ability to manipulate target biomolecules at user-defined time points, a central limitation is that most require overexpression of a protein of interest (POI) or its functional domain. Such overexpression frequently perturbs endogenous signaling pathways and cellular behaviors in a non-physiological manner, complicating investigation of cellular mechanisms under native conditions^4^.

Overcoming these limitations requires approaches that directly manipulate endogenous proteins without the need to overexpress the target protein itself^5^. Several approaches combining controllable domains with specific protein binders, such as intrabodies, have successfully demonstrated the inducible manipulation of endogenous proteins^6–8^. However, these tools have limited applicability because the repertoire of specific and robust protein binders remains small, and developing new binders is both challenging and time-consuming^9^. Furthermore, the structural and functional diversity of proteins makes it difficult to develop binder-dependent approaches into a single generalizable platform.

To overcome the need for target-specific binders and instead manipulate a wide range of endogenous proteins with a single platform, we focused on the application of synthetic biomolecular condensates. Biomolecular condensation driven by liquid-liquid phase separation (LLPS) is a common cellular mechanism for modulating protein activity^10–12^. Biomolecular condensates can sequester specific targets to inhibit them via steric hindrance, accumulate them to facilitate enzymatic processes, or release them to enable conditional activation. Building on this understanding, inducible platforms exploiting synthetic biomolecular condensation have been developed as simple and versatile strategies for regulating target protein activity^13–21^. While most of the existing techniques are designed for controlling exogenous proteins, a synthetic biomolecular condensate platform demonstrating inducible manipulation of endogenous mammalian proteins with downstream cellular functions remains elusive. By combining condensate scaffolds with genomic tagging or CRISPR-mediated knock-ins, inducible sequestration and release of endogenous proteins have been demonstrated in yeast using thermally responsive coiled-coil interactions^13^, and in *Drosophila* using optogenetic clustering based on Magnet photodimerizers^21^. However, the yeast system relies on temperature-dependent coiled-coil dissociation for client release, while the Magnet-based optogenetic system requires pre-incubation at 28°C for optimal dimerization. Both complicate direct application in mammalian cells maintained at 37°C. More recently, synthetic protein-recruiting/releasing condensate (SPREC) has been developed to sequester and release target proteins in mammalian cells and has been combined with a ligand-modulated anti-GFP nanobody^22^ (^GFP^LAMA) to target endogenous GFP-tagged ERK2^16, 23^. However, release efficiency for endogenous ERK2 remains relatively low (∼40%), and the functional consequences of this manipulation on downstream cellular behaviors have not been demonstrated. Therefore, a platform capable of sequestering and releasing endogenous proteins with high efficiency, and demonstrable downstream functional effects, remains an important unmet need.

For efficient sequestration and release of endogenous proteins, which are typically expressed at nanomolar-to-micromolar concentrations, it is crucial to employ a controllable affinity pair that exhibits low-nanomolar affinity yet can be effectively switched to near-zero affinity at a user-defined time point. Streptavidin binds an engineered streptavidin-binding peptide (SBP) with high affinity (*K*_d_ ∼ 2.5 nM), an interaction that can be rapidly outcompeted by biotin, whose extraordinary affinity for streptavidin (*K*_d_ ∼ 10 fM) drives displacement of SBP^24, 25^. The streptavidin-SBP system and the engineered Strep-Tactin system have previously been used to control overexpressed proteins for synchronizing secretory cargo trafficking (RUSH system)^26^, repositioning organelles^27^, and regulating protease activity^28^ in mammalian cells. While recent work combining the RUSH system with an endogenously SBP-tagged transferrin receptor (TfR) has enabled the study of endogenous TfR trafficking^29^, this system utilizes streptavidin localized in the endoplasmic reticulum or Golgi and is therefore suited to observing cargo transport in the secretory pathway rather than inhibiting protein function. In this study, we developed an engineered streptavidin condensate system for the inducible control of endogenous proteins in mammalian cells. We reasoned that anchoring streptavidin to synthetic biomolecular condensates would enable the tight compartmentalization and functional inhibition of endogenously SBP-tagged proteins, followed by their rapid release upon the addition of biotin (**Fig. 1a**). By genomically tagging endogenous proteins with SBP via CRISPR knock-in editing, we demonstrated that this engineered streptavidin condensate system can target a variety of essential cellular machinery, including molecular motors and the actin cytoskeleton, to control intracellular vesicle trafficking and lamellipodia formation. To achieve full temporal control over both sequestration and release, we further developed a dual-inducible system in which rapamycin triggers streptavidin condensation and POI sequestration, while subsequent biotin treatment drives rapid release, enabling precise temporal manipulation of endogenous protein activity.

**Figure 1.**
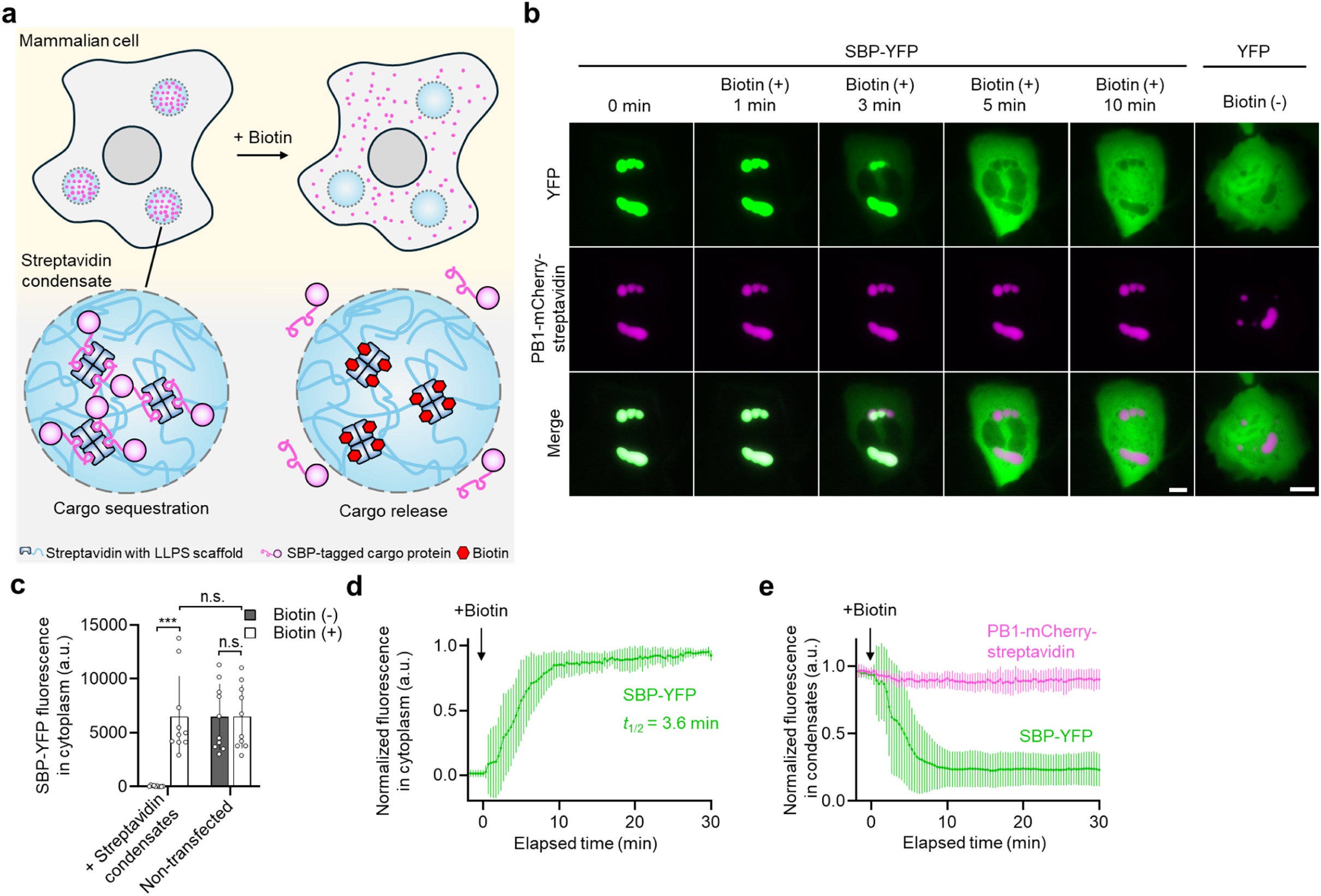
Streptavidin condensates allow biotin-inducible control of a target protein in mammalian cells. (**a**) Schematic of the streptavidin condensate system. Streptavidin fused with tandem oligomerizing or disordered domains forms biomolecular condensates within cells, enabling the containment and subsequent release of target cargo proteins tagged with streptavidin-binding peptide (SBP). (**b**) Time-lapse images of U2OS cells expressing streptavidin condensates encoded by PB1-mCherry-streptavidin (magenta) and SBP-YFP or control YFP (green). The YFP constructs used and the time point of biotin treatment are indicated. Scale bar: 10 µm. (**c**) Quantification of cytoplasmic SBP-YFP fluorescence from **b** before (grey) and after 10 min of biotin treatment (white). U2OS cells were transfected with or without PB1-mCherry-streptavidin as indicated. Dots represent individual data points and bars indicate mean ± s.d. (n = 10 cells from two independent experiments). *** *P* < 0.005; n.s., *P* > 0.05. (**d**) Kinetics of SBP-YFP in the cytoplasm. (**e**) Kinetics of SBP-YFP (green) and PB1-mCherry-streptavidin (magenta) within the condensates. Biotin was added at time 0. Lines and error bars indicate mean ± s.d. (n = 11 cells from two independent experiments).

## Results

### Sequestering and releasing target proteins by engineered streptavidin condensates

We fused several established LLPS scaffolds with streptavidin and co-expressed each with an SBP-fused GFP or HaloTag as a model cargo protein in human U2OS cells (**Supplementary Fig. 1**). All tested LLPS scaffolds (the PB1 domain fused with tetrameric AzamiGreen^30, 31^ or with mCherry (relying on streptavidin tetramerization for multivalency), tandem RGG domains^32^, and Y15 peptide fused with AzamiGreen^33^) successfully formed condensates with streptavidin and sequestered SBP-tagged cargo proteins. We further confirmed that biotin treatment induced the release of the sequestered cargos. These results demonstrated that trapping and releasing target proteins via synthetic streptavidin condensates is compatible with a range of LLPS scaffolds. Because of its efficient sequestration and negligible leakage (**Supplementary Fig. 1c**), we focused on streptavidin fused to the PB1 domain for subsequent experiments. SBP is a 38-amino-acid peptide; a shortened 24-amino-acid variant has been reported to retain similar affinity for streptavidin *in vitro*^24^. We verified that YFP fused to the shorter SBP variant was efficiently trapped in and subsequently released from the synthetic streptavidin condensates to the same extent as full-length SBP; thus, we utilized the shorter SBP variant in the following experiments (**Supplementary Fig. 2**). Time-lapse live imaging confirmed that PB1-mCherry-streptavidin condensates efficiently captured SBP-YFP with no observable cytosolic leakage, while control YFP lacking SBP was unaffected (**Fig. 1b–d and Supplementary Video 1**). Upon addition of biotin, the sequestered SBP-YFP was rapidly released from the condensates, with its cytoplasmic concentration recovering to levels comparable to cells lacking condensates within minutes (*t*_1/2_ = 3.6 min) (**Fig. 1b–e and Supplementary Video 1**). We also demonstrated that this system can trap and release SBP-tagged proteins across several commonly used mammalian cell lines, including HEK293T, HeLa, and COS-7 (**Supplementary Fig. 3**). Lastly, we directly compared this system with the recently developed ^GFP^LAMA-SPREC system^23^, which can trap GFP-tagged proteins and release them in response to trimethoprim (TMP) (**Supplementary Fig. 4a**). We confirmed that PB1-MontiRed-^GFP^LAMA can trap cytosolic GFP-tagged cargo proteins but with some leakage (**Supplementary Fig. 4b**). In addition, while streptavidin condensates can release most sequestered proteins upon biotin treatment, the TMP-induced cargo release from PB1-MontiRed-^GFP^LAMA was relatively inefficient (**Supplementary Fig. 4c**), consistent with a previous report^23^. These results establish a streptavidin condensate system capable of sequestering and releasing target proteins in a rapid and chemically inducible manner within mammalian cells.

### Manipulation of endogenous motor proteins and intracellular trafficking in mammalian cells

Having established robust control of exogenous cargo, we asked whether the streptavidin condensate system could be applied to manipulate endogenous proteins in mammalian cells. KIF5B is the most abundant kinesin-1 family member in most cell types and drives plus-end-directed transport along microtubules^34, 35^. KIF5B transports a broad range of cargos, including proteins, mRNAs, vesicles, and organelles, making it a central mediator of intracellular trafficking. To manipulate endogenous KIF5B with the streptavidin condensate, we generated human near-diploid HT-1080 cells homozygously tagged with SBP at the endogenous *KIF5B* alleles (**Fig. 2a, b**). Using a CRISPR-mediated knock-in approach and subsequent single-cell isolation via cell sorting^36, 37^, we successfully established an HT-1080 homozygous clone featuring SBP-YFP at the N-terminus of endogenous KIF5B (*KIF5B^SBP/SBP^*), which was confirmed by genomic PCR (**Fig. 2c**), sequencing (**Fig. 2d**), and Western blot (**Fig. 2e**).

**Figure 2.**
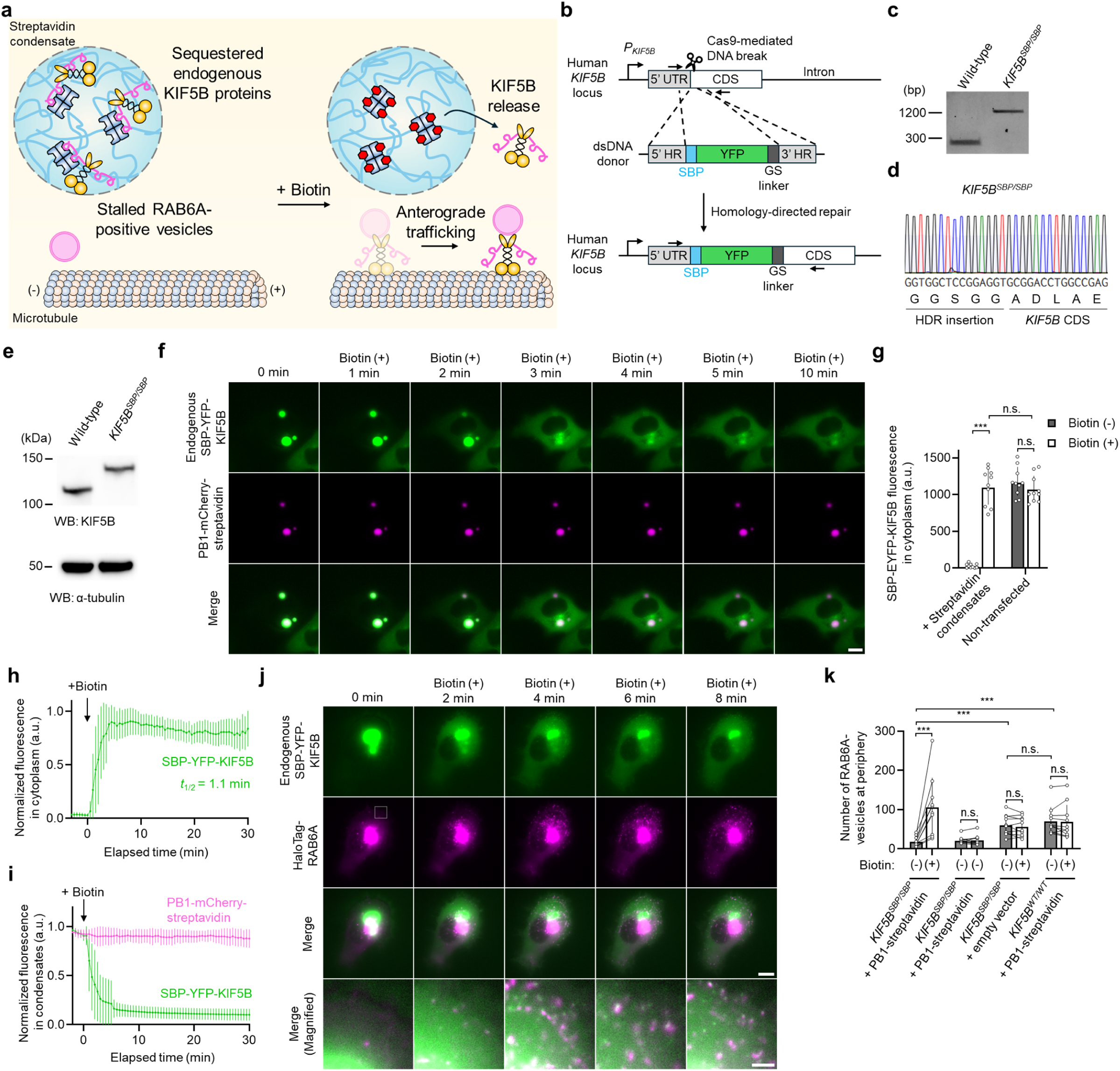
Manipulation of endogenous kinesin KIF5B and KIF5B-mediated RAB6A-positive vesicle trafficking by streptavidin condensates. (**a**) Schematics of endogenous KIF5B manipulation using streptavidin condensates. Endogenous *KIF5B* alleles in HT-1080 cells are homozygously tagged with SBP-YFP at the N-terminus via CRISPR-mediated knock-in editing (*KIF5B^SBP/SBP^*). The SBP-YFP-tagged KIF5B is sequestered and inhibited within streptavidin condensates, halting RAB6A-positive vesicle movement. KIF5B containment is released by biotin treatment, restoring KIF5B-mediated RAB6A vesicle trafficking. (**b**) Genome editing strategy of SBP-YFP knock-in in human *KIF5B* locus. Primers for genotyping are designed to flank the integration site. (**c**) Genomic PCR for genotyping *KIF5B^SBP/SBP^*cells. (**d**) Sequence confirmation of SBP-YFP integration in *KIF5B^SBP/SBP^*cells. (**e**) Western blot for endogenous KIF5B in wild-type and *KIF5B^SBP/SBP^*cells. (**f**) Time-lapse images of *KIF5B^SBP/SBP^* cells expressing streptavidin condensates encoded by PB1-mCherry-streptavidin (magenta). Endogenous SBP-YFP-KIF5B is shown in green. The timing of biotin treatment is indicated. Scale bar: 10 µm. (**g**) Quantification of SBP-YFP-KIF5B in the cytoplasm before (grey) and after 5 min of biotin treatment (white). *KIF5B^SBP/SBP^* cells were transfected with or without the streptavidin condensate construct as indicated. Dots are individual data points and bars indicate mean ± s.d. (n = 10 cells from three independent experiments). (**h**) Kinetics of endogenous SBP-YFP-KIF5B in the cytoplasm. (**i**) Kinetics of endogenous SBP-YFP-KIF5B (green) and PB1-mCherry-streptavidin (magenta) in the condensates. Biotin was added at time 0. Lines and error bars indicate mean ± s.d. (n = 10 cells from three independent experiments). (**j**) Live imaging of RAB6A-positive vesicles during the manipulation of endogenous KIF5B by streptavidin condensates. *KIF5B^SBP/SBP^* cells were transfected with PB1-streptavidin and HaloTag-RAB6A (magenta). Magnified merged images of the regions outlined by white squares are also shown. Scale bar: 10 µm (main images) and 2 µm (insets). (**k**) The number of RAB6A-positive vesicles at the cell periphery before (grey) and after 7.5 min of biotin treatment (white). *KIF5B^SBP/SBP^* or wild-type (*KIF5B^WT/WT^*) cells were transfected with PB1-streptavidin or an empty vector as indicated. Paired dots are individual data points and bars indicate mean ± s.d. (n = 10 cells from two independent experiments). *** *P* < 0.005; n.s., *P* > 0.05.

To test whether the streptavidin condensate could trap and release endogenously tagged KIF5B, we transfected *KIF5B^SBP/SBP^* cells with PB1-mCherry-streptavidin and conducted live-cell imaging (**Fig. 2f, Supplementary Video 2**). Endogenous KIF5B was efficiently sequestered into streptavidin condensates with no observable cytosolic leakage and released upon biotin treatment with 90% release efficiency (**Fig. 2f, g**). We quantified the kinetics of KIF5B release, confirming rapid release (*t*_1/2_ = 1.1 min) (**Fig. 2h, i**).

Following this successful manipulation of endogenous KIF5B in mammalian cells, we set out to test whether this system could control KIF5B-mediated intracellular trafficking. Among various cargos trafficked by KIF5B, we focused on RAB6A-positive vesicles, which are Golgi-derived exocytotic vesicles transported to the plasma membrane by KIF5B activity (**Fig. 2a**)^38, 39^. *KIF5B^SBP/SBP^* cells were transfected with PB1-streptavidin and HaloTag-RAB6A and subjected to live-cell imaging (**Fig. 2j, Supplementary Video 3**). In cells where endogenous KIF5B was sequestered within the condensates, we observed a significant depletion of RAB6A-positive vesicles at the cell periphery compared to non-transfected cells (**Fig. 2k**). Upon biotin treatment, KIF5B was released into the cytoplasm, localized to RAB6A vesicles within minutes, and transported them toward the cell periphery. These results demonstrate that the streptavidin condensate can trap and release endogenous KIF5B to control intracellular trafficking.

We subsequently applied the streptavidin condensate to control a different class of endogenous motor proteins. Cytoplasmic dynein is a minus-end-directed microtubule motor that drives retrograde transport of diverse cargos from the cell periphery toward the cell center^40^. During endocytosis, internalized receptors and ligands are sorted into RAB5A-positive early endosomes, which mature into RAB7A-positive late endosomes^41–44^. Both early and late endosomes depend on dynein-mediated transport for their centripetal positioning, which is essential for efficient cargo sorting.

To test whether the streptavidin condensate could control endogenous dynein activity and dynein-mediated early endosome localization, we tagged the endogenous cytoplasmic dynein heavy chain (encoded by *DYNC1H1*) with SBP-mNeonGreen (mNG) using a CRISPR knock-in approach (**Fig. 3a**). We verified the successful generation of homozygous SBP-mNG-DYNC1H1 HT-1080 cells (*DYNC1H1^SBP/SBP^*) by genomic PCR, sequencing, and Western blot (**Supplementary Fig. 5a–c**). Overexpression of the streptavidin condensate construct in *DYNC1H1^SBP/SBP^*cells efficiently trapped endogenous SBP-mNG-DYNC1H1, which could then be rapidly released upon biotin treatment (*t*_1/2_ = 1.7 min) (**Fig. 3b–d, Supplementary Video 4**). These results show that the streptavidin condensate can trap exceptionally large proteins (∼530 kDa), underscoring the versatility of the system. We visualized early endosomes by co-expressing HaloTag-RAB5A and confirmed that, in the absence of biotin, RAB5A-positive vesicles accumulated heavily at the cell periphery compared to cells lacking the streptavidin condensates (**Fig. 3b, e**). Notably, the accumulation of RAB5A-positive vesicles was often concentrated at focal peripheral sites rather than uniformly distributed along the cell periphery. This pattern may reflect the polarized nature of endocytic activity in migrating cells, which is concentrated at discrete hotspots such as leading edges^45^. After inducing dynein release with biotin, the peripheral RAB5A-positive vesicles began moving toward the cell center, and the number of RAB5A vesicles at the cell periphery decreased. This demonstrated that the streptavidin condensate can trap dynein to inhibit its retrograde transport function and subsequently release it to transport stalled peripheral RAB5A vesicles to the cell center. Next, we visualized late endosomes by expressing HaloTag-RAB7A. In cells where dynein was trapped within the condensates, RAB7A vesicles accumulated more prominently at the cell periphery compared to cells without condensates and wild-type cells (**Fig. 3f, g, Supplementary Video 5**). Following biotin treatment, the RAB7A vesicles translocated to the cell center, reducing the number of RAB7A-positive endosomes at the cell periphery (**Fig. 3g**). These results demonstrate that the streptavidin condensate can effectively manipulate endogenous motor proteins to control intracellular transport.

**Figure 3.**
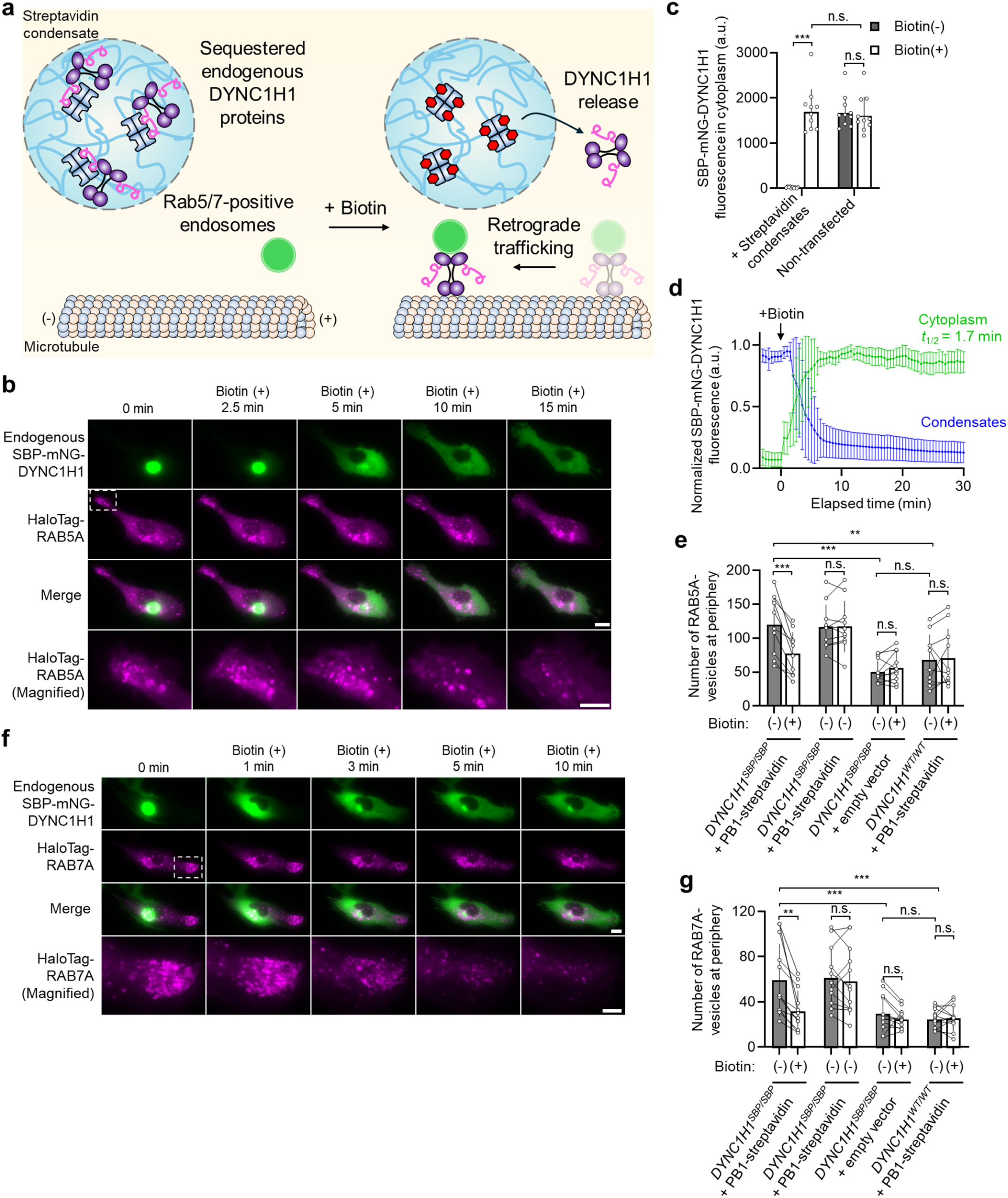
Manipulation of endogenous cytoplasmic dynein and dynein-mediated early and late endosome trafficking using streptavidin condensates. (**a**) Schematics of endogenous cytoplasmic dynein manipulation using streptavidin condensates. Endogenous *DYNC1H1* alleles in HT-1080 cells are homozygously tagged with SBP-mNeonGreen (mNG) at the N-terminus via CRISPR-mediated knock-in editing (*DYNC1H1^SBP/SBP^*). The SBP-mNG-tagged DYNC1H1 is sequestered and inhibited within streptavidin condensates, disrupting retrograde movement of early and late endosomes marked by RAB5A and RAB7A, respectively. DYNC1H1 containment is released by biotin treatment, restoring cytoplasmic dynein-mediated early and late endosomal trafficking. (**b**) Time-lapse images of *DYNC1H1^SBP/SBP^* cells expressing streptavidin condensates encoded by PB1-streptavidin and HaloTag-RAB5A (magenta). Endogenous SBP-mNG-DYNC1H1 is shown in green. Magnified images of the regions outlined by white squares are also shown. The timing of biotin treatment is indicated. Scale bar: 10 µm (main images) and 5 µm (insets). (**c**) Quantification of SBP-mNG-DYNC1H1 in the cytoplasm before (grey) and after 8 min of biotin treatment (white). *DYNC1H1^SBP/SBP^*cells were transfected with or without the streptavidin condensate construct as indicated. Dots are individual data points and bars indicate mean ± s.d. (n = 10 cells from two independent experiments). (**d**) Kinetics of endogenous SBP-mNG-DYNC1H1 in the cytoplasm (green) and condensates (blue). Biotin was added at time 0. Lines and error bars indicate mean ± s.d. (n = 10 cells from three independent experiments). (**e**) The number of RAB5A-positive vesicles at the cell periphery before (grey) and after 10 min of biotin treatment (white). *DYNC1H1^SBP/SBP^* or wild-type (*DYNC1H1^WT/WT^*) cells were transfected with PB1-streptavidin or an empty vector as indicated. Paired dots are individual data points and bars indicate mean ± s.d. (n = 10 cells from two independent experiments). (**f**) Live imaging of RAB7A-positive vesicles during the manipulation of endogenous DYNC1H1 by streptavidin condensates. *DYNC1H1^SBP/SBP^* cells were transfected with PB1-streptavidin and HaloTag-RAB7A (magenta). Magnified images of the regions outlined by white squares are also shown. Scale bar: 10 µm (main images) and 5 µm (insets). (**g**) The number of RAB7A-positive endosomes at the cell periphery before (grey) and after 10 min of biotin treatment (white). *DYNC1H1^SBP/SBP^*or *DYNC1H1^WT/WT^* cells were transfected with PB1-streptavidin or an empty vector as indicated. Paired dots are individual data points and bars indicate mean ± s.d. (n = 12 cells from two independent experiments). ** *P* < 0.01; *** *P* < 0.005; n.s., *P* > 0.05.

### Manipulation of actin dynamics by controlling endogenous Arp2/3 complexes in mammalian cells

To further demonstrate the broad applicability of the streptavidin condensate, we adapted the system to manipulate the endogenous Arp2/3 complex, an essential regulator of actin filament dynamics (**Fig. 4a)**^46^. Branched actin networks generate the mechanical forces required for membrane remodeling during processes such as cell migration, exocytosis, and phagocytosis. These networks are nucleated by the Arp2/3 complex, which binds pre-existing actin filaments and initiates the growth of new branch filaments. The Arp2/3 complex consists of seven subunits: ARP2, ARP3, and ARPC1–5. To establish inducible control of the endogenous Arp2/3 complex, we tagged the C-terminus of endogenous ARPC3 with mScarlet-SBP in HT-1080 cells (*ARPC3^SBP/SBP^*), as previous studies have shown that endogenous tagging at this site preserves Arp2/3 complex function^47^. Successful homozygous integration at the expected *ARPC3* alleles was verified by genomic PCR, sequencing, and Western blot (**Supplementary Fig. 5d–f**). In contrast to *KIF5B^SBP/SBP^* and *DYNC1H1^SBP/SBP^*(**Figs. 2**, **3**), *ARPC3^SBP/SBP^* showed significantly lower ARPC3 abundance compared to wild-type (**Supplementary Fig. 5f**). Despite reduced ARPC3 abundance detected by Western blot, immunofluorescence using the anti-ARPC3 antibody revealed that *ARPC3^SBP/SBP^* cells displayed membrane ruffles with ARPC3 localization comparable to wild-type (**Supplementary Fig. 5g, h**). The mScarlet signal in *ARPC3^SBP/SBP^* cells also colocalized with these ruffles, confirming that the endogenously tagged ARPC3-mScarlet-SBP localizes to active sites of actin-driven membrane remodeling. These results indicate that the Arp2/3 complex containing endogenously tagged ARPC3-mScarlet-SBP remains functional in driving actin-mediated membrane protrusions. Thus, we set out to test whether streptavidin condensates can manipulate endogenously tagged ARPC3 functions in *ARPC3^SBP/SBP^* cells.

**Figure 4.**
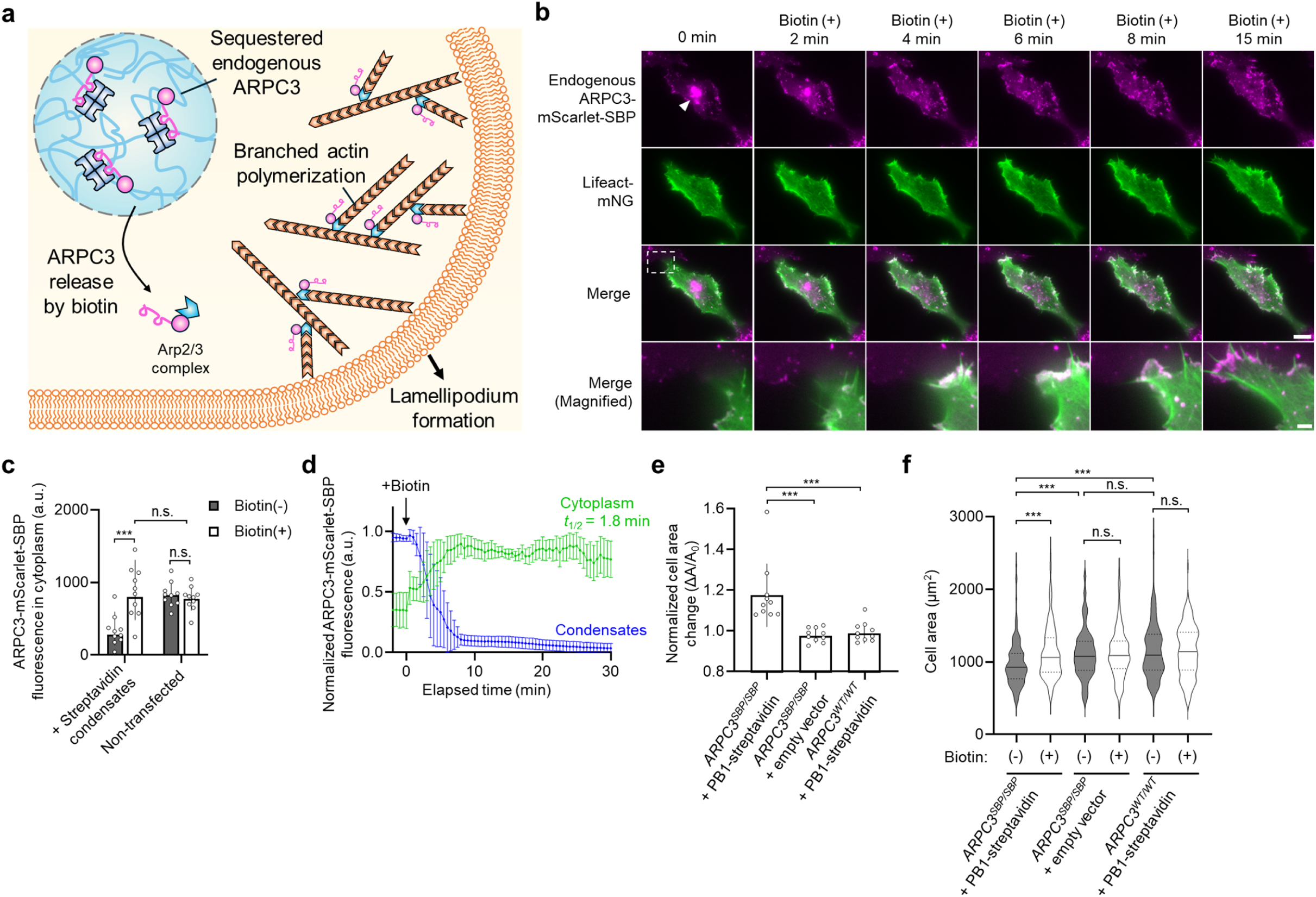
Manipulation of endogenous Arp2/3 complex and Arp2/3-mediated branched actin polymerization using streptavidin condensates. (**a**) Schematics of endogenous Arp2/3 complex manipulation using streptavidin condensates. Endogenous *ARPC3* alleles in HT-1080 cells are homozygously tagged with mScarlet-SBP at the C-terminus via CRISPR-mediated knock-in editing (*ARPC3^SBP/SBP^*). The mScarlet-SBP-tagged ARPC3 is sequestered into streptavidin condensates, resulting in inefficient Arp2/3-mediated cell protrusion and cell area reduction. ARPC3 containment is released by biotin treatment, restoring branched actin polymerization and cell area expansion. (**b**) Live imaging of cellular morphology with F-actin marker, Lifeact-mNG (green) during the manipulation of endogenous ARPC3-mScarlet-SBP (magenta) by streptavidin condensates. *ARPC3^SBP/SBP^*cells were transfected with PB1-streptavidin and Lifeact-mNG. A white arrowhead indicates trapped ARPC3 before biotin treatment. Magnified merged images of the regions outlined by white squares are also shown. The timing of biotin treatment is indicated. Scale bar: 10 µm (main images) and 2 µm (insets). (**c**) Quantification of ARPC3-mScarlet-SBP in the cytoplasm before (grey) and after 5 min of biotin treatment (white). *ARPC3^SBP/SBP^* cells were transfected with or without the streptavidin condensate construct as indicated. Dots are individual data points and bars indicate mean ± s.d. (n = 10 cells from three independent experiments). (**d**) Kinetics of endogenous ARPC3-mScarlet-SBP in the cytoplasm (green) and condensates (blue). Biotin was added at time 0. Lines and error bars indicate mean ± s.d. (n = 10 cells from three independent experiments). (**e**) Normalized cell area changes in **b** by biotin treatment. *ARPC3^SBP/SBP^* or wild-type (*ARPC3^WT/WT^*) cells were transfected with PB1-streptavidin or an empty vector as indicated. Dots are individual data points and bars indicate mean ± s.d. (n = 10 cells from three independent experiments). (**f**) Quantification of cell area with (white) and without (grey) biotin treatment. *ARPC3^SBP/SBP^* cells were transfected with PB1-streptavidin or an empty vector as indicated. HaloTag-CAAX was co-transfected as the membrane marker for cell segmentation. Lines show the median, and dotted lines indicate the first and third quartiles from three independent experiments (n = 160, 155, 178, 144, 159, and 161 cells, from left to right). *** *P* < 0.005; n.s., *P* > 0.05.

To assess the functional consequences of endogenous ARPC3 manipulation, *ARPC3^SBP/SBP^* cells were co-transfected with the streptavidin condensate construct and Lifeact-mNeonGreen to visualize actin polymerization^48^. We confirmed that approximately 60% of endogenously tagged ARPC3-mScarlet-SBP was sequestered in the condensates and rapidly released into the cytoplasm following biotin treatment (*t*_1/2_ = 1.8 min) (**Fig. 4b–d, Supplementary Video 6**). The release of endogenous ARPC3 facilitated lamellipodia formation marked by both ARPC3 and Lifeact and drove cellular area expansion (**Fig. 4b, e**). To more quantitatively assess whether ARPC3 sequestration reduced Arp2/3-driven cell area expansion and whether its release restored this expansion, we conducted high-throughput analysis of cell area using the Cellpose-SAM model^49^. This confirmed an approximately 15% reduction in cell area upon ARPC3 sequestration and its recovery following biotin-induced release (**Fig. 4f**). These results demonstrate that the streptavidin condensate can manipulate endogenous Arp2/3 to control membrane dynamics.

### Inducible sequestration and release of endogenously tagged proteins in mammalian cells

To further extend the controllability of endogenous proteins, we combined biotin-triggered release with inducible streptavidin condensate formation to create a dual-inducible trap-and-release system (**Fig. 5a**). To achieve inducible condensation of streptavidin, we integrated a rapamycin-inducible dimerization system, in which FKBP and FRB form a heterodimer in the presence of rapamycin^1^. Because multivalent interactions drive LLPS, we designed PB1-FKBP and streptavidin-mScarlet-FRB constructs to increase inter-scaffold multivalency in an inducible manner^30^. While both PB1-FKBP and streptavidin-mScarlet-FRB form independent multimers, they do not spontaneously form phase-separated condensates (**Fig. 5b, Supplementary Video 7**). However, inducing FKBP-FRB dimerization via rapamycin treatment enhances these multivalent interactions, driving the formation of biomolecular condensates (**Fig. 5c**). We confirmed that the generated condensates effectively trapped over 90% of cytosolic SBP-YFP after rapamycin treatment (**Fig. 5d**). Upon subsequent biotin treatment, the sequestered SBP-YFP was rapidly released within minutes, recovering its cytoplasmic concentration to pre-rapamycin levels.

**Figure 5.**
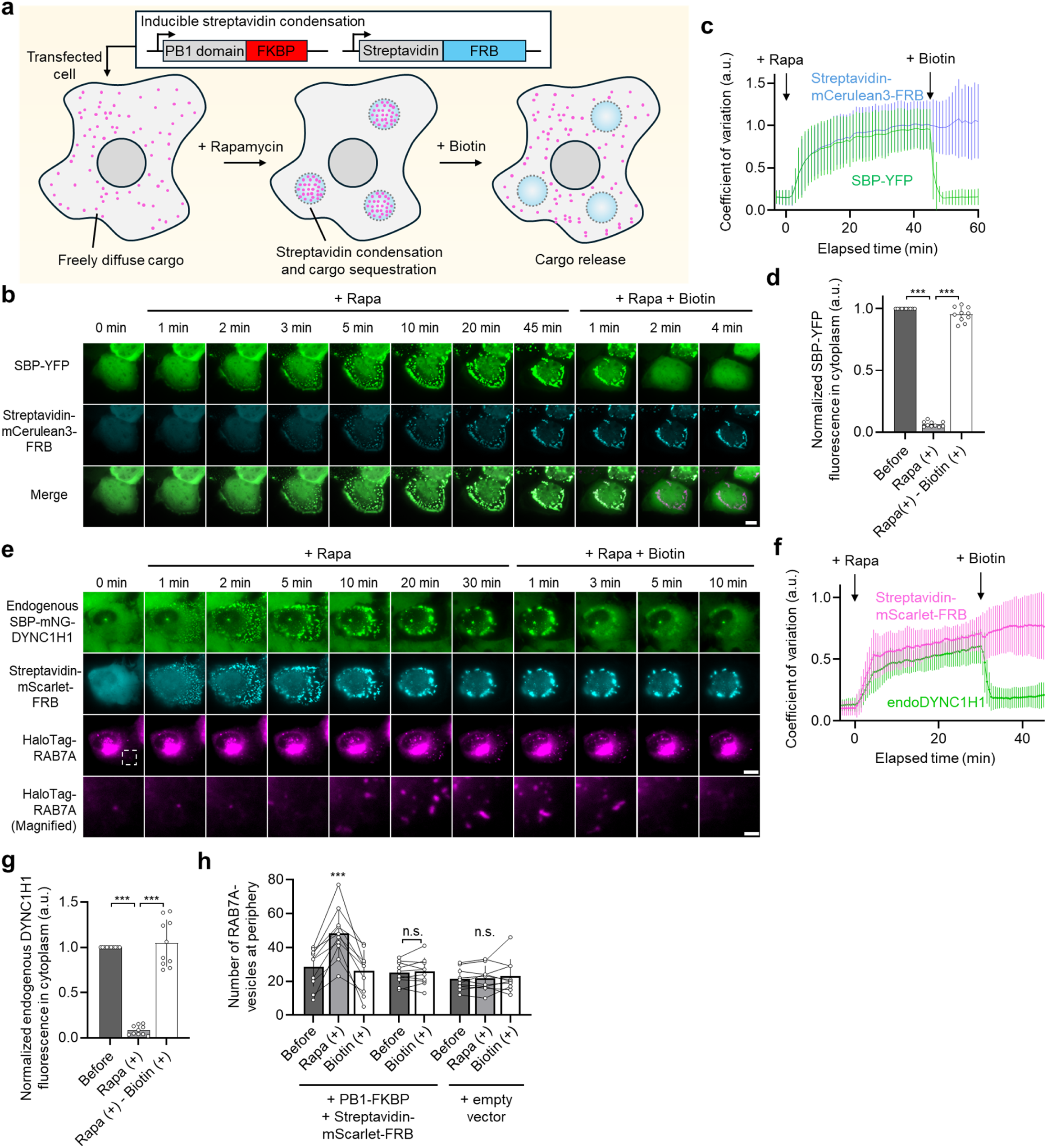
Controllable sequestration and release of target endogenous proteins using rapamycin-induced streptavidin condensation. (**a**) Schematics of rapamycin-induced streptavidin condensation for sequestration and subsequent release of a target protein in an inducible manner. Streptavidin fused with FRB interacts with SBP-tagged target cargo proteins. Upon rapamycin treatment, the FRB forms a heterodimer with FKBP fused with a PB1 multimerizing domain, resulting in the formation of streptavidin condensates that sequester the SBP-tagged protein. (**b**) Time-lapse images of U2OS cells expressing the rapamycin-induced streptavidin condensation system encoded by PB1-FKBP and streptavidin-mCerulean3-FRB (cyan) and SBP-YFP (green). The timing of rapamycin and biotin treatment is indicated. Scale bar: 10 µm. (**c**) Kinetics of standard deviations of integrated intensity density distributions of SBP-YFP (green) and streptavidin-mCerulean3-FRB (blue) in **b**. Rapamycin and biotin were added at time 0 and 45, respectively. Lines and error bars indicate mean ± s.d. (n = 14 cells from two independent experiments). (**d**) Quantification of cytoplasmic SBP-YFP in **b** before (dark grey) and after rapamycin (light grey) and subsequent biotin treatment (white). Dots are individual data points and bars indicate mean ± s.d. (n = 10 cells from three independent experiments). (**e**) Time-lapse images of *DYNC1H1^SBP/SBP^* cells expressing the rapamycin-induced streptavidin condensation system encoded by PB1-FKBP and streptavidin-mScarlet-FRB (cyan) with HaloTag-RAB7A (magenta). The timing of rapamycin and biotin treatment is indicated. Magnified images of the regions outlined by white squares are also shown. Scale bar: 10 µm (main images) and 2 µm (insets). (**f**) Kinetics of standard deviations of integrated intensity density distributions of endogenous SBP-mNG-DYNC1H1 (green) and streptavidin-mScarlet-FRB (magenta) in **e**. Rapamycin and biotin were added at time 0 and 30, respectively. Lines and error bars indicate mean ± s.d. (n = 10 cells from three independent experiments). (**g**) Quantification of cytoplasmic SBP-mNG-DYNC1H1 in **e** before (dark grey) and after rapamycin (light grey) and subsequent biotin treatment (white). Dots are individual data points and bars indicate mean ± s.d. (n = 10 cells from three independent experiments). (**h**) The number of RAB7A-positive endosomes at the cell periphery before (dark grey) and after rapamycin (light grey) and subsequent biotin treatment (white). *DYNC1H1^SBP/SBP^* cells were transfected with PB1-FKBP-P2A-streptavidin-mScarlet-FRB or an empty vector as indicated. Paired dots are individual data points and bars indicate mean ± s.d. (n = 10 cells from three independent experiments). *** *P* < 0.005; n.s., *P* > 0.05.

To demonstrate that this dual-inducible system could control endogenous targets, we introduced it into the *DYNC1H1^SBP/SBP^* cells (**Fig. 3a**). We confirmed that this system enables the rapid sequestration of endogenous DYNC1H1 into rapamycin-induced streptavidin condensates, followed by its release into the cytosol upon biotin treatment (**Fig. 5e, f, Supplementary Video 8**). We quantified the endogenous DYNC1H1 intensity in the cytoplasm before and after rapamycin and subsequent biotin treatment, confirming the sequestration of over 90% of endogenous DYNC1H1 and its efficient release (**Fig. 5g**). To determine whether this system could sequentially control dynein-mediated cellular behavior, we monitored the distribution of RAB7A-marked late endosomes (**Fig. 5e**). Prior to treatment, RAB7A-positive vesicles accumulated near the cell center. Following the rapamycin-induced trapping of the endogenous cytoplasmic dynein heavy chain, the RAB7A vesicles displayed a more dispersed distribution, reflected by an increased number at the cell periphery (**Fig. 5h**). Upon biotin-induced release of sequestered DYNC1H1, RAB7A vesicles relocated from the cell periphery back to the cell center, resulting in a decrease in peripheral vesicle number. Collectively, these results demonstrate that the dual-inducible system enables chemically controlled capture and release of endogenous proteins in living mammalian cells.

## Discussion

In this study, we developed a streptavidin condensate platform for the chemically inducible control of endogenous proteins in mammalian cells. By using this platform with CRISPR-mediated knock-in of a short SBP tag into a target locus, we demonstrated tight sequestration and rapid release of target endogenous proteins across diverse cellular targets, including microtubule motor proteins KIF5B and DYNC1H1 and the Arp2/3 complex subunit ARPC3. We further confirmed that the sequestration and release of target proteins affect the related downstream cellular functions such as vesicle trafficking and actin-driven membrane dynamics, demonstrating that our system achieves functional manipulation.

A key advantage of this platform is that it targets endogenous proteins directly, avoiding the non-physiological stoichiometry introduced by overexpression-based tools. Unlike intrabody-based approaches, which require a unique high-affinity binder for each new target, our system requires only insertion of the universal short SBP tag via a well-established gene editing strategy, making the platform readily extensible to virtually any cytosolic protein^50^. We directly compared our system to the recently reported ^GFP^LAMA-SPREC condensate system, which demonstrated trap and release of endogenous GFP-tagged ERK2 in mammalian cells^23^. While ^GFP^LAMA-SPREC shows only ∼40% release efficiency as originally reported^23^, our platform achieves near-complete release of cargo proteins, underscoring its advantage. Our platform also complements well-established degradation-based systems such as AID^51^ or dTAG^52^, which remove proteins over timescales of tens of minutes to hours. In contrast, our system provides inducible reactivation: it liberates sequestered proteins and restores their functions within minutes (*t*_1/2_ = 1.1 min for KIF5B, 1.7 min for DYNC1H1, 1.8 min for ARPC3), making the two strategies suited to different experimental questions. Because both the efficiency and kinetics of cargo release depend on the fluidity of protein condensates, we envision that these properties could be tuned and improved by integrating insights from rational engineering of synthetic condensates with specific mechanical properties^53, 54^. Broader design principles for engineered condensates have been reviewed recently^55^.

Despite these strengths, several limitations warrant consideration. First, as a general limitation in knock-in gene editing experiments, endogenous tagging may affect protein expression and function. In our experiments, knock-in tagging outcomes varied across target genes, with *KIF5B* and *DYNC1H1* expression preserved and *ARPC3* reduced; nonetheless, the residual tagged ARPC3 retained function and was sufficient to demonstrate inducible control of Arp2/3-mediated membrane dynamics. Possible explanations of reduced expression by endogenous tagging include altered mRNA and protein stability, and steric hindrance by an introduced exogenous tag. To minimize these possibilities, the insertion sites, linker sequences, and choice of fluorescent tags (if used) should be carefully considered. Second, because the streptavidin condensates form in the cytosol, the platform is currently best suited to cytosolic proteins; membrane-associated, nuclear, or organelle-resident proteins may be physically inaccessible to condensate-mediated sequestration. As recent work has demonstrated that synthetic condensates can be engineered to form in the nucleus and on the cytoplasmic surfaces of organelle membranes^14, 56^, future work should explore whether targeting streptavidin condensate formation to specific subcellular compartments can overcome this limitation. Third, because biotin is inherently cell-impermeable and its uptake by mammalian cells relies on the ubiquitously expressed sodium multivitamin transporter, the onset of cargo release may vary across experimental conditions and cell types, as previously reported^57^. Recent advances in cell-permeable biotin analogs should lower this barrier^58^. Lastly, while the strong affinity between streptavidin and biotin provides highly efficient cargo release from streptavidin condensates, it makes this system irreversible. However, the engineered M88 streptavidin mutant demonstrates the possibility of achieving reversible streptavidin-biotin binding^59, 60^; M88 employs an engineered disulfide bond in a critical loop near the binding pocket, and it releases biotinylated ligands effectively under reducing conditions. Thus, by integrating chemically sensitive streptavidin into synthetic condensates, truly reversible cargo manipulation could be achieved.

In summary, the streptavidin condensate platform provides a rapid, scalable, and target-agnostic approach for the inducible control of endogenous proteins in mammalian cells. Its compatibility with diverse targets, multiple cell lines, and a dual-inducible configuration positions it as a broadly useful tool for dissecting spatiotemporal protein regulation. We anticipate that this platform will find wide application in basic cell biological research and may prove valuable in biotechnological applications requiring precise, chemically controlled manipulation of specific endogenous proteins.

## Acknowledgements

We thank Jack Gregory, Ian Llaneza, and Arshadul Hak for participating in initial preliminary experiments. We thank Dr. Takanari Inoue for providing the opportunity to conduct preliminary experiments in his laboratory. This work was supported by National Institutes of Health (R35GM159972, to Y. N.) and start-up funds from Ohio State University. This work was also supported by The Ohio State University Comprehensive Cancer Center and the National Institutes of Health under grant number P30CA016058. We thank the Flow Cytometry Shared Resource and the Genomics Shared Resource at The Ohio State University Comprehensive Cancer Center, Columbus, OH for sorting the knock-in cells and DNA sequencing, respectively.

## Author contributions

T. K., C. J. W., I. L. and Y. N. designed, performed, and analyzed experiments. T. K. and Y. N. wrote the manuscript. All authors edited the manuscript. Y. N. supervised the project.

## Methods

### Plasmid construction

All plasmids encoding chimeric proteins in this study were generated by standard seamless In-Fusion Cloning (Takara Bio, 638949). All PCR reactions for plasmid construction were carried out with Q5 High-Fidelity DNA Polymerase (NEB, M0491L) following the manufacturer’s protocol. For streptavidin condensate constructs, a cDNA encoding streptavidin was amplified from Halo-VSVG-RUSH (Addgene plasmid #166907; a gift from Jennifer Lippincott-Schwartz). cDNAs encoding PB1 domain and AzamiGreen were amplified from pCMV-PB1-AG-FRB (Addgene plasmid #178862; a gift from Shinya Tsukiji). A cDNA encoding RGG-RFP-RGG was amplified from pcDNA_RGG-RFP-RGG (Addgene plasmid #124934; a gift from Matthew Good, Daniel Hammer, and Benjamin Schuster). A cDNA encoding mScarlet was amplified from pmScarlet_peroxisome_C1 (Addgene plasmid #85063; a gift from Dorus Gadella). A cDNA encoding MontiRed was amplified from pCAGGS-PB1-MontiRed-LOV2dark (Addgene plasmid #178874; a gift from Shinya Tsukiji). A cDNA encoding ^GFP^LAMA was amplified from pCDNA3.0_mitoLAMA-F98 (Addgene plasmid #130704; a gift from Kai Johnsson). A cDNA encoding RAB6A was amplified from EGFP-Rab6A (Addgene plasmid #49469; a gift from Marci Scidmore). cDNAs encoding GFP, YFP, mCherry, mCerulean3, FKBP, FRB, RAB5A and RAB7A were gifts from Takanari Inoue. Sequences encoding streptavidin binding peptide (MDEKTTGWRGGHVVEGLAGELEQLRARLEHHPQGQREP), the short variant of SBP (GHVVEGLAGELEQLRARLEHHPQG), Y15 peptide (YEYKYEYKYEYKYEY), Lifeact (MGVADLIKKFESISKEE), CAAX motif from KRAS (NPPDESGPGCMSCKCVLS), and P2A peptide (GSGATNFSLLKQAGDVEENPGP) were included as primer extensions. These PCR fragments were cloned into pcDNA3.1(+) vector digested with *Nhe*I and *Eco*RI. To construct the SBP-GFP virus vector, the amplified fragment was cloned into FUGW vector (a gift from Takanari Inoue) digested with *Age*I and *Eco*RI. To construct the HaloTag-RAB6A virus vector, the amplified fragment was cloned into pSLQ1658-dCas9-EGFP vector (Addgene plasmid #51023; a gift from Bo Huang and Stanley Qi) digested with *Bgl*II and *Eco*RI. All plasmids were verified by Sanger sequencing or Nanopore whole plasmid sequencing. The primary structures of the streptavidin condensates and SBP-fused cargo proteins are described in **Supplementary Note 1**.

### Cell culture

U2OS (ATCC, HTB-96), HeLa (ATCC, CCL-2), COS-7 (ATCC, CRL-1651), HEK293T (ATCC, CRL-3216) and HT-1080 (Millipore Sigma, 85111505-1VL) cells were maintained in DMEM (Corning, 10-013-CV) supplemented with 10% FBS (Corning, 35-010-CV), 1% Pen-Strep (Thermo Fisher, 15140163) at 37 °C under 5% CO_2_. These cell lines have been authenticated by manufacturers and checked for mycoplasma contamination prior to use in experiments in this paper.

### Donor DNA preparation

Double-strand DNA donors encoding SBP-YFP, SBP-mNeonGreen and mScarlet-SBP with 40 bp homology arms were generated by standard PCR. The sequences of the DNA donors used in this study are described in **Supplementary Note 2**. Plasmids encoding SBP-YFP, SBP-mNeonGreen or mScarlet-SBP were used as a PCR template. PCR primers targeting these sequences with homology arms were synthesized without chemical modification (IDT). PCR amplicons were generated with Q5 High-Fidelity DNA polymerase (NEB, M0491L), following the manufacturer’s protocol. The amplicons were purified by MinElute PCR Purification Kit (Qiagen, 28004) and resuspended in ddH_2_O to 1 µg/µL.

### Generation of human cell lines with endogenous protein tagging

crRNAs targeting human *KIF5B*, *DYNC1H1*, and *ARPC3* were synthesized by Integrated DNA Technologies (IDT). The target sequences of the crRNAs are described in **Supplementary Table 1**. The crRNAs were mixed with tracrRNAs (IDT) at a final concentration of 50 µM, denatured at 95 °C for 5 min to anneal, and cooled to room temperature. To form a Cas9:guide RNA complex, 2 µl of the crRNA:tracrRNA mixture was mixed with 1 µl of purified Cas9 nuclease at 61 µM (IDT) and incubated for 15 minutes at room temperature. The Cas9:guide RNA complex was introduced into HT-1080 cells using 4D-Nucleofector system (Lonza) with the SF Cell Line 4D-Nucleofector X kit (Lonza, V4XC-2032) and FF-113 program. Cells were plated on 6-well culture plates (Corning), and collected for single cell sorting 7 days after electroporation. The top 5% of fluorescent cells were sorted into 96-well culture plates by BD FACSAria III Cell Sorter (BD) at the flow cytometry shared resource at the Ohio State University Comprehensive Cancer Center. The isolated cells were grown with changing medium every 3 days. After 15 days of growth, the cells were split into two separate culture plates for genotyping and further cell expansion. To identify clones with homozygous integration of SBP and fluorescent proteins, genomic PCR, sequencing, and Western blotting or immunofluorescence were performed.

### Genomic PCR

Genomic DNA was extracted from single-cell clones using a QuickExtract DNA Extraction Solution (Lucigen, QE09050) according to the manufacturer’s protocol. To verify homozygous integration of SBP and fluorescent protein tags at the target locus, genomic PCR was performed using primers flanking the insertion site. For human *KIF5B* and *ARPC3*, touchdown PCR was carried out with Q5 High-Fidelity DNA Polymerase under the following conditions: initial denaturation at 98 °C for 30 s, followed by first 10 cycles of 98 °C for 10 s, 72-62 °C, - 1 °C/cycle for 20 s, and 72 °C for 30 s and second 25 cycles of 98 °C for 10 s, 62 °C for 20 s, and 72 °C for 30 s, and a final extension at 72 °C for 2 min. For human DYNC1H1, 3-step PCR was carried out with PrimeSTAR GXL DNA polymerase (Takara Bio, R050A) under the following conditions: initial denaturation at 98 °C for 10 s, followed by 30 cycles of 98 °C for 10 s, 60 °C for 15 s, and 68 °C for 60 s, and a final extension at 68°C for 2 min. The primer sequences used for genotyping are listed in **Supplementary Table 2**. PCR products were resolved on a 2% agarose gel and visualized with Midori Green Xtra (Bulldog Bio, MG10). Successful knock-in was confirmed by the expected shift in amplicon size compared to wild-type alleles, and homozygosity was verified by the absence of a wild-type-sized band.

### Western blot

Cells were lysed in sample buffer (50 mM Tris-HCl, 2% SDS, and 10% glycerol) supplemented with cOmplete protease inhibitor cocktail (Roche, 11697498001). Lysates were clarified by centrifugation at 12,000 × g for 10 min at 4 °C, and protein concentrations were determined using a BCA Protein Assay Kit (Thermo Scientific, A55865). Equal amounts of protein (10 µg per lane) were separated on 4–15% Mini-PROTEAN TGX Precast Protein Gels (Bio-Rad, 4561085) and transferred to PVDF membranes (Bio-Rad, 1620177). Membranes were blocked with 4% BSA in TBST (Cell Signaling Technology, 9997) for 1 h at room temperature, then incubated with primary antibodies overnight at 4 °C. The following primary antibodies were used: anti-KIF5B (Cell Signaling Technology, 62696, 1:1,000), anti-DYNC1H1 (Proteintech, 12345-1-AP, 1:2,000), anti-ARPC3 (BD, 612234, 1:1,000) and anti-tubulin (Proteintech, 11224-1-AP, 1:10,000) as a loading control. After washing three times with TBST, membranes were incubated with HRP-conjugated anti-rabbit secondary antibodies (Cell Signaling Technology, 7074S, 1:1,000) or anti-mouse secondary antibodies (Cell Signaling Technology, 7076S, 1:1,000) for 1 h at room temperature. Bands were detected using SignalFire™ ECL Reagent (Cell Signaling Technology, 6883) and imaged on ChemiDoc MP system (Bio-Rad). Band sizes were compared against Precision Plus Protein Kaleidoscope Prestained Protein Standards (Bio-Rad, 1610375).

### Immunofluorescence

Cells were seeded at a density of 10,000 cells per well into a fibronectin-coated 8-well glass-bottom chamber (Cellvis, C8-1.5H-N). After 24 h incubation, cells were washed with PBS (−) three times and fixed with 4% paraformaldehyde in PBS (−) for 10 min at room temperature. Next, cells were washed with PBS (−) three times and permeabilized and blocked with 2% BSA and 0.1% Triton X-100 in PBS (−) for 30 min at room temperature. After blocking, cells were incubated with primary antibody against ARPC3 (BD, 612234, 1:500) diluted in blocking buffer overnight at 4 °C. Cells were then washed with PBS (−) three times and incubated with diluted anti-mouse secondary antibody conjugated with Alexa Fluor 647 (Thermo Fisher Scientific, A-31571, 1:1,000) and 14 nM of Acti-stain™488 fluorescent phalloidin (Cytoskeleton, PHDG1-A) in blocking buffer for 1 h at room temperature. Cells were washed with PBS (−) three times again and subjected to fluorescence imaging performed on an Eclipse Ti inverted fluorescence microscope (Nikon) equipped with 60× oil-immersion objective lens and ORCA-Fusion BT Digital sCMOS camera (Hamamatsu Photonics).

### Virus production

For HaloTag-RAB6A virus, GP2-293 packaging cell lines (Takara Bio, 631458) were seeded at 6 × 10^6^ cells in a 10-cm culture dish (Corning) with 10 ml of complete DMEM and incubated for 24 h at 37 °C in 5% CO_2_. Transfer plasmid and envelope plasmid (pMD2.G, Addgene #12259; a gift from Didier Trono) were transfected using PEI MAX (Polysciences, 24765) following the standard protocol. For SBP-GFP lentivirus, HEK293T cells were seeded instead and transfected with transfer plasmid, pMD2.G and psPAX2 packaging plasmid (psPAX2, Addgene #12260; a gift from Didier Trono). At 24 h post transfection, the medium was replaced with fresh DMEM. The viral supernatant was collected at 48 and 72 h post transfection and concentrated by a standard polyethylene glycol precipitation protocol.

### Generation of stable cell lines

For generating wild-type HT-1080 cells and *KIF5B^SBP/SBP^*cells stably expressing HaloTag-RAB6A, as well as U2OS cells stably expressing SBP-GFP, cells were seeded at a density of 40,000 cells/well into 6-well plates (Corning). The next day, the cells were transduced with the virus. The transduction was performed with serum-free plain DMEM with 6 µg/ml of polybrene. 48 h after transduction, antibiotic screening was performed using 2 µg/ml puromycin. The puromycin screening was completed for 7 days and the expression was confirmed by live-cell imaging.

### Live-cell imaging

For wide-field fluorescence and total internal reflection fluorescence (TIRF) imaging in this paper, an Eclipse Ti inverted fluorescence microscope equipped with 60× oil-immersion objective lens and ORCA-Fusion BT Digital sCMOS camera was used. Time-lapse imaging was conducted at 37 °C, 5% CO_2_ and more than 90% humidity using a stage-top incubator (Tokai Hit).

For initial screening experiments and characterization during development of the streptavidin condensate, U2OS, HeLa, COS-7, and HEK293T cells were seeded at a density of 40,000 cells/well in 400 µL of complete medium into an 8-well glass-bottom chamber. Immediately after cell plating, the cells were transfected using 0.6 µg of PEI MAX (Polysciences, 24765) and 0.2 µg of DNA plasmids as indicated in the figure legends. After 24 h incubation, the cells were subjected to wide-field fluorescence imaging at 1-min intervals for 30 min. For induction of the release of SBP-tagged model proteins, cells were treated with 200 µM biotin (Sigma-Aldrich, B4639). For kinetic analysis of SBP-YFP release, time-lapse wide-field imaging was conducted 3 min prior to the addition of 200 µM biotin and continued for 30 min post-addition. Images were acquired at 20-sec intervals throughout the experiment.

For the endogenous SBP-tagged protein release experiment, wild-type HT-1080 cells, *KIF5B^SBP/SBP^* cells, *DYNC1H1^SBP/SBP^*cells, and *ARPC3^SBP/SBP^* cells were nucleofected with indicated DNA plasmids by 4D-Nucleofector system (Lonza) using the SF Cell line 4D-Nucleofector X Kit S (Lonza) and the FF-113 program. Then, the nucleofected cells were seeded at a density of 60,000 (*KIF5B^SBP/SBP^*), 80,000 (*DYNC1H1^SBP/SBP^*), and 120,000 (*ARPC3^SBP/SBP^*) cells/well into an 8-well glass-bottom chamber with fibronectin coating (Sigma-Aldrich, F1114). After 24 h incubation, the cells were incubated with 50 nM of Janelia Fluor 646 HaloTag Ligand (Promega, GA1120) for 15 min for visualizing HaloTag-fused markers. Time-lapse fluorescence imaging was conducted 3 min prior to the addition of 2 mM biotin and continued for 30 min post-addition. Images were acquired at 30-sec intervals throughout the experiment. For *ARPC3^SBP/SBP^*cells with Lifeact-mNG, images were acquired at 1-min intervals.

For kinetic analyses of SBP-YFP with rapamycin-inducible streptavidin condensation, U2OS cells were seeded at a density of 40,000 cells/well into an 8-well glass-bottom chamber. Right after cell plating, the cells were co-transfected with indicated DNA plasmids using PEI MAX as described above. Time-lapse wide-field fluorescence imaging was conducted 3 min before the addition of 100 nM rapamycin and continued for 45 min post-addition of rapamycin. Then, 200 µM biotin was added to the cell medium and the acquisition proceeded for an additional 15 minutes. Images were acquired at 20-sec intervals.

For live-cell imaging of *DYNC1H1^SBP/SBP^* cells using the rapamycin-induced trapping system, indicated DNA plasmids were nucleofected into the cells by 4D-Nucleofector system (Lonza) as described above. Cells were then seeded at a density of 80,000 cells/well into a fibronectin-coated 8-well glass-bottom chamber. After incubation of the cells for 24 h, the cells were incubated with 50 nM of Janelia Fluor 646 HaloTag Ligand for 15 min. Time-lapse TIRF imaging was performed at 30-sec intervals, beginning 3 min prior to the addition of 500 nM rapamycin and continuing for 30 min post-addition. Then, 2 mM biotin was added to the cell medium and the acquisition proceeded for an additional 15 minutes.

### Image analysis

For quantification of SBP-tagged cargo sequestration and release, three ROIs were manually drawn in the cytoplasmic region (neither in the nucleus nor in the condensates) for each cell and the mean fluorescence intensities of SBP-tagged cargos were then measured from the ROIs at indicated time points. For quantification of overexpressed SBP-fused protein release kinetics, we segmented the boundaries of intracellular condensates from the PB1-mCherry-streptavidin signal in the mCherry (mCh) channel using the Make Binary command in ImageJ/FIJI (NIH). To subtract background fluorescence intensity, the maximum fluorescence intensities of YFP and mCh from a region of interest (ROI) outside the cell were measured. After background subtraction from each pixel, the mean fluorescence intensities of YFP and mCh within each delineated condensate region were measured. For quantification of SBP-YFP release kinetics in the cytoplasm, the condensate binary masks were expanded using the Erode command in ImageJ/FIJI to exclude signal bleeding from the condensates into the surrounding cytoplasm. The mean fluorescence intensities of YFP were then measured across the whole-cell region excluding the expanded condensate mask.

For quantification of endogenously tagged KIF5B, DYNC1H1, and ARPC3 release kinetics, the fluorescence intensities of the tagged endogenous proteins and condensate scaffolds were individually measured in condensate and cytoplasmic regions as described above for overexpressed SBP-fused proteins in U2OS cells. To compensate for photobleaching, mean cytoplasmic fluorescence intensities from endogenous proteins at each time point were divided by the mean intensities of control cells (in which tagged endogenous proteins exhibited normal cytoplasmic localization) imaged in parallel at the corresponding time points.

For quantification of RAB-positive vesicles, the cell periphery was defined as follows. ROIs were drawn to encompass each single cell by manual tracing, then shrunk by 6 µm (for RAB6A) or 4 µm (for RAB5A and RAB7A) using the Enlarge command in ImageJ/FIJI with a negative length value. The cellular area outside the contracted ROIs was defined as the cell periphery. RAB-positive vesicles in the cell periphery were then manually counted at the first frame (biotin(−)) and after 7.5 min (RAB6A) or 10 min (RAB5A, RAB7A) of biotin treatment (biotin(+)).

For quantification of cell area expansion during endogenous ARPC3 manipulation using streptavidin condensates, we segmented the boundaries of cells from the Lifeact-mNG channel using the Make Binary command in ImageJ/FIJI. The areas of the segmented cells were then measured at the first frame (CA_biotin(−)_) and after 30 min of biotin treatment (CA_biotin(+)_). The normalized cell area change was calculated as CA_biotin(+)_/CA_biotin(−)_. To segment cells and quantify absolute cell areas, fluorescent images of HaloTag-CAAX from cells were processed using Cellpose-SAM model^49^ (https://github.com/MouseLand/cellpose). The segmentation by Cellpose was converted to ROIs and separately confirmed the expression of streptavidin condensates. The selected ROIs were then imported into ImageJ for quantifying cell area.

For quantification of rapamycin-induced condensate formation and sequestration of SBP-tagged proteins, after background subtraction as described above, an ROI was manually drawn in the cytoplasm for each cell as follows: the size of the ROI was from 100 to 200 µm^2^ (approximately 2.0 × 10^4^ to 4.0 × 10^4^ pixels); the ROI was placed within the cytoplasmic region excluding the nucleus throughout the time-lapse. The standard deviation of fluorescence intensities within each ROI was measured at each time point using ImageJ/FIJI. For quantification of SBP-tagged cargo sequestration and release by rapamycin-induced streptavidin condensation system, three ROIs were manually drawn in the cytoplasmic region for each cell at the first frame (before), after 45 min of rapamycin treatment (rapa (+)), and after further 5 min of biotin treatment (rapa (+) - biotin (+)). For quantification of RAB7A-positive vesicles during reversible DYNC1H1 control, the cell periphery was defined as described above. RAB7A-positive vesicles in the cell periphery were then manually counted at the first frame (before), after 30 min of rapamycin treatment (rapa (+)), and after further 30 min of biotin treatment (rapa (+) - biotin (+)).

### Reproducibility and statistics

Pairwise comparisons were conducted using the Wilcoxon signed-rank test, Friedman test, and Brunner-Munzel test in GraphPad Prism 10 (GraphPad Software) or Microsoft Excel (Microsoft). No sample size estimations were performed, and our sample sizes are consistent with those typically used in live-cell imaging experiments. For live-cell experiments, we selected cells as follows: to ensure efficient sequestration, we manually selected cells expressing streptavidin condensates with no observable cytosolic leakage of cargo proteins before biotin treatment. The ARPC3 release experiment was an exception; for the ARPC3 release experiment, we selected the cells showing at least 50% intensity reduction in the cytoplasm by condensate-mediated sequestration. No randomization was used because the study does not involve allocation into different experimental groups. No blinding was used because samples were not grouped and randomized. For all experiments in this manuscript, at least two independent experiments were performed and all attempts at replication were successful. For confirmation of the homozygous knock-in editing by Western blot, the representative blot image was selected from two independent experiments. The numbers of independent experiments and analyzed cells are indicated in each figure legend.

## Data availability

The datasets generated during this study are available from the corresponding author upon request. The key plasmids in this paper will be deposited at Addgene prior to publication.

