## Supporting Information for "An engineered streptavidin condensate platform for chemically inducible control of endogenous proteins in mammalian cells"

**Supplementary Figure 1. Streptavidin fused with diverse multimerizing domains and intrinsically disordered domains forms biomolecular condensates that enable biotin-inducible cargo release in mammalian cells.** (**a**–**d**) Representative images of U2OS cells expressing PB1-AzamiGreen-streptavidin (**a**), Y15-AzamiGreen-streptavidin (**b**), PB1-mCherry-streptavidin (**c**), and streptavidin-RGG-mCherry-RGG (**d**). Model cargo proteins, SBP-HaloTag (**a, b**) or SBP-YFP (**c, d**) are also co-transfected. Scale bar: 10 µm. (right) Quantification of cytoplasmic SBP-tagged cargo intensity before (grey) and after 30 min of biotin treatment (White) is shown. Paired dots are individual data points and bars indicate mean ± s.d. (n = 10 cells from two independent experiments). *** $P < 0.005$; n.s., $P > 0.05$.

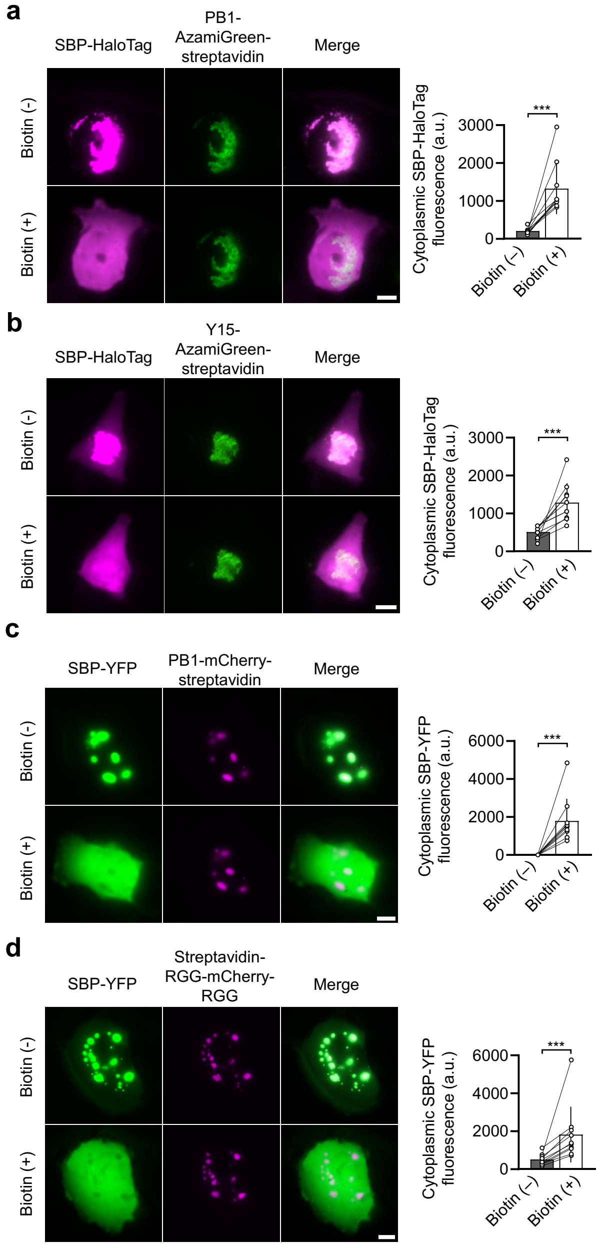

**Supplementary Figure 2. A shorter variant of streptavidin-binding peptide can be used for efficient sequestration of cargo proteins into streptavidin condensates.** (**a**) Representative images of U2OS cells expressing PB1-mCherry-streptavidin (magenta) and YFP fused with SBP ($SBP_{36aa}$) or a shorter SBP variant ($SBP_{24aa}$) (green). The SBP-YFP constructs used and the presence of biotin treatment are indicated. Scale bar: 10 μm. Representative images from two individual experiments are shown. (**b**) Quantification of cytoplasmic SBP-YFP intensity in **a** before (grey) and after biotin treatment (white). The SBP-YFP constructs used are indicated. Dots are individual data points and bars indicate mean ± s.d. (n = 10 cells from two independent experiments). *** $P < 0.005$; n.s., $P > 0.05$.

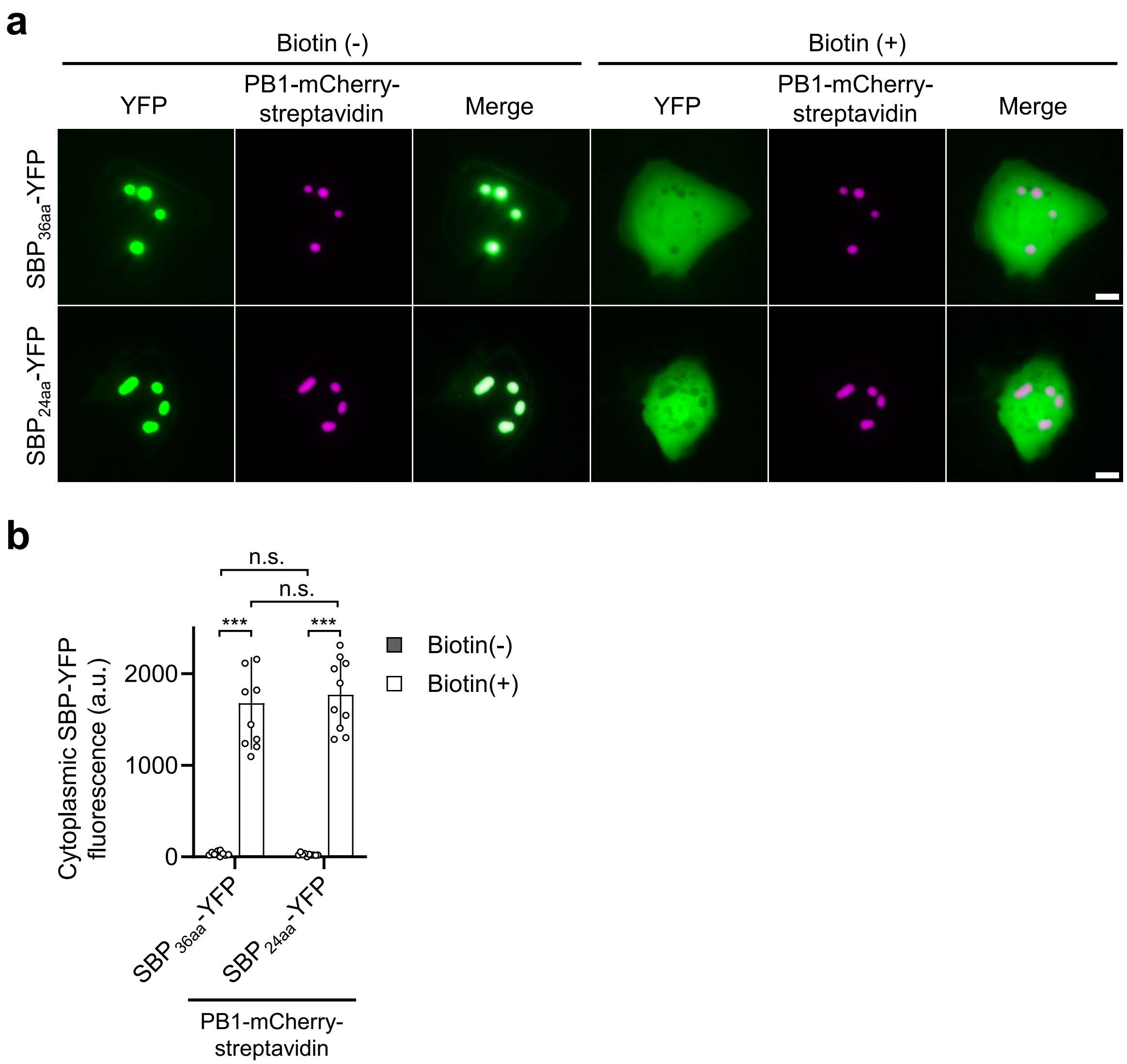

**Supplementary Figure 3. Streptavidin condensates-mediated sequestration and release of cargo proteins in commonly used mammalian cell lines.** (**a**) Representative images of COS-7, HEK293T, and HeLa cells expressing PB1-mCherry-streptavidin (magenta) and SBP-YFP (green). The cell lines used and the presence of biotin treatment are indicated. Scale bar: 10 μm. Representative images from two individual experiments are shown. (**b**) Quantification of cytoplasmic SBP-YFP intensity in **a** before (grey) and after (white) biotin treatment. The cell lines used are indicated. Dots are individual data points and bars indicate mean ± s.d. (n = 10 cells from two independent experiments). *** $P < 0.005$; n.s., $P > 0.05$.

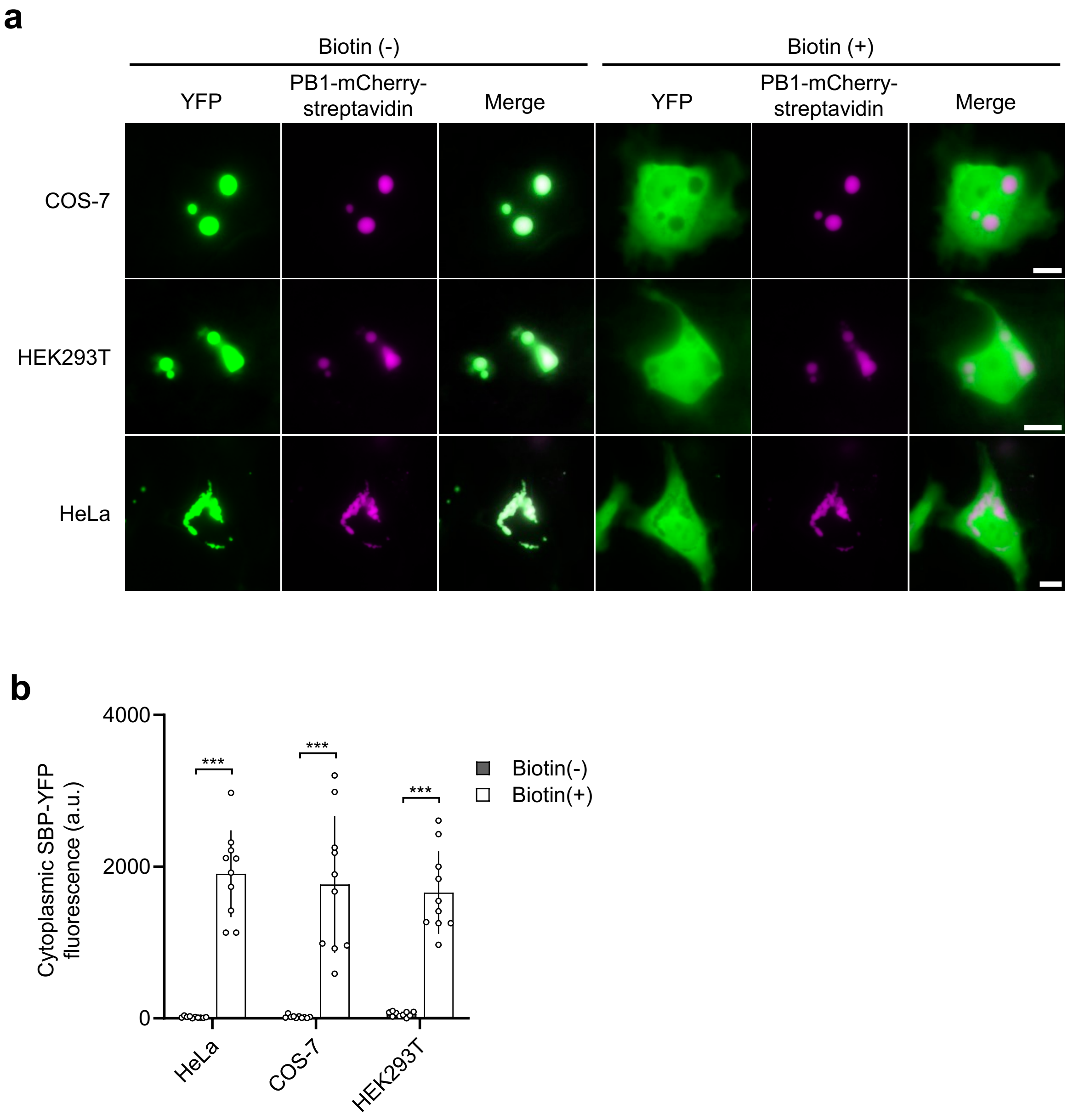

**Supplementary Figure 4. Side-by-side comparison between streptavidin condensate and $^{GFP}$LAMA-based condensate.** (**a**) Representative images of U2OS cells expressing SBP-GFP (green) and PB1-mCherry-streptavidin or PB1-MontiRed-$^{GFP}$LAMA (magenta). Cells were treated with biotin (for PB1-mCherry-streptavidin) or trimethoprim (for PB1-MontiRed-$^{GFP}$LAMA). Scale bar: 10 μm. (**b**) Quantification of cytoplasmic SBP-GFP intensity in **a** before (grey) and after 15 min of drug treatment (white). Dots are individual data points and bars indicate mean ± s.d. (n = 10 cells from two individual experiments). (**c**) Relative SBP-GFP fluorescence change within condensates upon drug treatment. Dots are individual data points and bars indicate mean ± s.d. (n = 10 cells from two individual experiments). ** $P < 0.01$; *** $P < 0.005$; n.s., $P > 0.05$.

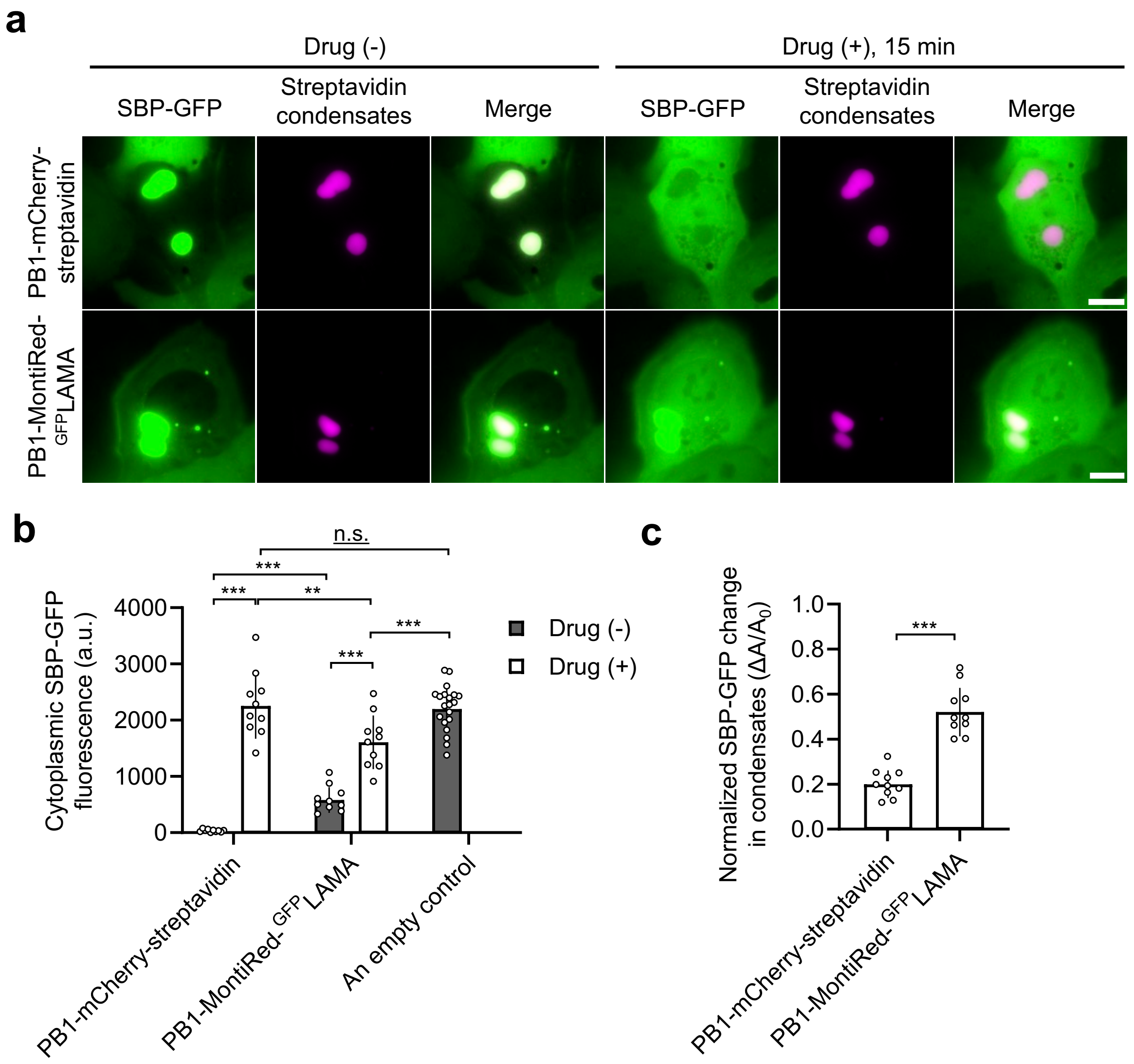

**Supplementary Figure 5. Validation of homozygously knock-in HT-1080 cells.** (**a**) Genomic PCR for genotyping *DYNC1H1*$^{SBP/SBP}$ cells. (**b**) Sequencing confirmation of *DYNC1H1*$^{SBP/SBP}$ knock-in. (**c**) Western blot for endogenous DYNC1H1. (**d**) Genomic PCR for genotyping *ARPC3*$^{SBP/SBP}$ cells. (**e**) Sequencing confirmation of *ARPC3*$^{SBP/SBP}$. (**f**) Western blot for endogenous ARPC3. (**g, h**) Immunofluorescence of wild-type (**g**) and *ARPC3*$^{SBP/SBP}$ (**h**) cells for F-actin (green) and endogenous ARPC3 (magenta). Endogenously tagged ARPC3-mScarlet-SBP in *ARPC3*$^{SBP/SBP}$ is shown in red. Scale bar: 10 µm (main images) and 2 µm (insets).

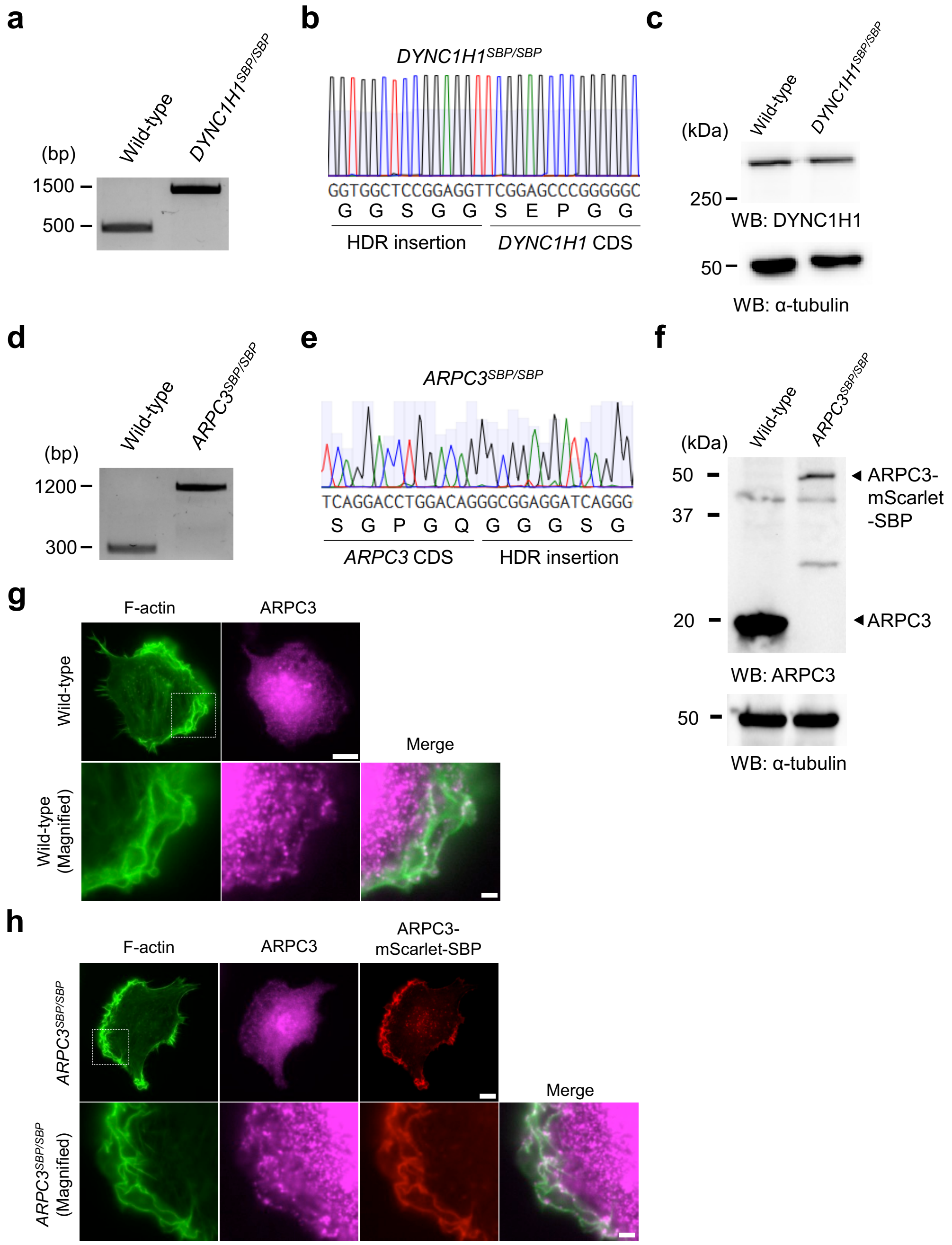

**Supplementary Table 1|** Target sequences of crRNAs used in the paper.

| Target gene | Target sequence (5' to 3') |
|---|---|
| Human *KIF5B* | GGCCCCGGCTGCGAGAAAGA |
| Human *DYNC1H1* | TCCCGAGCGCGACACCATGT |
| Human *ARPC3* | CCGGGCTCCCTTCACTGTCC |

**Supplementary Table 2|** Genomic PCR primers for genotyping CRISPR knock-in clones.

| Target gene | Forward primer (5' to 3') | Reverse primer (5' to 3') |
|---|---|---|
| Human *KIF5B* | GCCATGATGGATCGGAAGTG | ATCACGACCGTGTCTTCTCC |
| Human *DYNC1H1* | TGTCTCTTGCTGGCTGTCTC | CATCCAGGGGTCATCTGCAG |
| Human *ARPC3* | TGACCCCGCTGTTCATTCTG | CCACTTTCTTTTCTCCCACCC |

**Supplementary Note 1|** Amino acid sequences of the key constructs used in the paper.

>>SBP-YFP

MGHVVEGLAGELEQLRARLEHHPQGGGGGSGGGSGVSKGEELFTGVVPILVELDGDVN
GHKFSVSGEGEGDATYGKLTLKFICTTGKLPVPWPTLVTTFGYGLQCFARYPDHMKQHD
FFKSAMPEGYVQERTIFFKDDGNYKTRAEVKFEGDTLVNRIELKGIDFKEDGNILGHKLEY
NYNSHNVYIMADKQKNGIKVNFKIRHNIEDGSVQLADHYQQNTPIGDGPVLLPDNHYLSY
QSALSKDPNEKRDHMVLLEFVTAAGITLGMDEL

>>PB1-mCherry-streptavidin

MASLTVKAYLLGKEDAAREIRRFSFCCSPEPEAEAEAAAGPGPCERLLSRVAALFPALRP
GGFQAHYRDEDGDLVAFSSDEELTMAMSYVKDDIFRIYIKEKTGSGSGGSGAGGSAGSG
AGGSAGSGAGGSAGSGAGGSAGSGAGGSASGSSPGSGSGGSGAGGSAGSGAGGSA
GSGAGGSAGSGAGGSAGSGAGGSASGSSPGNSADGGGGSGGSGGSGGGSTQGGSM
VMVSKGEEDNMAIIKEFMRFKVHMEGSVNGHEFEIEGEGEGRPYEGTQTAKLKVTKGGP
LPFAWDILSPQFMYGSKAYVKHPADIPDYLKLSFPEGFKWERVMNFEDGGVVTVTQDSS
LQDGEFIYKVKLRGTNFPSDGPVMQKKTMGWEASSERMYPEDGALKGEIKQRLKLKDG
GHYDAEVKTTYKAKKPVQLPGAYNVNIKLDITSHNEDYTIVEQYERAEGRHSTGGMDELY
KGGGGSGGGSGDPSKDSKAQVSAAEAGITGTWYNQLGSTFIVTAGADGALTGTYESAV
GNAESRYVLTGRYDSAPATDGSGTALGWTVAWKNNYRNAHSATTWSGQYVGGAEARI
NTQWLLTSGTTEANAWKSTLVGHDTFTKVKPSAASIDAAKKAGVNNGNPLDAVQQ

>>PB1-mCerulean3-streptavidin

MASLTVKAYLLGKEDAAREIRRFSFCCSPEPEAEAEAAAGPGPCERLLSRVAALFPALRP
GGFQAHYRDEDGDLVAFSSDEELTMAMSYVKDDIFRIYIKEKTGSGSGGSGAGGSAGSG
AGGSAGSGAGGSAGSGAGGSAGSGAGGSASGSSPGSGSGGSGAGGSAGSGAGGSA
GSGAGGSAGSGAGGSAGSGAGGSASGSSPGNSADGGGGSGGSGGSGGGSTQGGSM
VVSKGEELFTGVVPILVELDGDVNGHKFSVSGEGEGDATYGKLTLKFICTTGKLPVPWPT
LVTTLSWGVQCFARYPDHMKQHDFFKSAMPEGYVQERTIFFKDDGNYKTRAEVKFEGDT
LVNRIELKGIDFKEDGNILGHKLEYNAIHGNVYITADKQKNGIKANFGLNCNIEDGSVQLAD
HYQQNTPIGDGPVLLPDNHYLSTQSKLSKDPNEKRDHMVLLEFVTAAGITLGMDELGGG
GSGGGSGDPSKDSKAQVSAAEAGITGTWYNQLGSTFIVTAGADGALTGTYESAVGNAES
RYVLTGRYDSAPATDGSGTALGWTVAWKNNYRNAHSATTWSGQYVGGAEARINTQWLL
TSGTTEANAWKSTLVGHDTFTKVKPSAASIDAAKKAGVNNGNPLDAVQQ

>>PB1-streptavidin

MASLTVKAYLLGKEDAAREIRRFSFCCSPEPEAEAEAAAGPGPCERLLSRVAALFPALRP
GGFQAHYRDEDGDLVAFSSDEELTMAMSYVKDDIFRIYIKEKTGSGSGGSGAGGSAGSG
AGGSAGSGAGGSAGSGAGGSAGSGAGGSASGSSPGSGSGGSGAGGSAGSGAGGSA
GSGAGGSAGSGAGGSAGSGAGGSASGSSPGNSADGGGGSGGSGGSGGGSTQGGSM
VGGGSGGGSGGDPSKDSKAQVSAAEAGITGTWYNQLGSTFIVTAGADGALTGTYESAV
GNAESRYVLTGRYDSAPATDGSGTALGWTVAWKNNYRNAHSATTWSGQYVGGAEARI
NTQWLLTSGTTEANAWKSTLVGHDTFTKVKPSAASIDAAKKAGVNNGNPLDAVQQ

>>PB1-FKBP

MASLTVKAYLLGKEDAAREIRRFSFCCSPEPEAEAEAAAGPGPCERLLSRVAALFPALRP
GGFQAHYRDEDGDLVAFSSDEELTMAMSYVKDDIFRIYIKEKTGSGSGGSGAGGSAGSG
AGGSAGSGAGGSAGSGAGGSAGSGAGGSASGSSPGSGSGGSGAGGSAGSGAGGSA
GSGAGGSAGSGAGGSAGSGAGGSASGSSPGNSADGGGGSGGSGGSGGGSTQGGSM
VGVQVETISPGDGRTFPKRGQTCVVHYTGMLEDGKKFDSSRDRNKPFKFMLGKQEVIRG
WEEGVAQMSVGQRAKLTISPDYAYGATGHPGIIPPHATLVFDVELLKLE

>>Streptavidin-mScarlet-FRB

MDPSKDSKAQVSAAEAGITGTWYNQLGSTFIVTAGADGALTGTYESAVGNAESRYVLTG
RYDSAPATDGSGTALGWTVAWKNNYRNAHSATTWSGQYVGGAEARINTQWLLTSGTTE
ANAWKSTLVGHDTFTKVKPSAASIDAAKKAGVNNGNPLDAVQQGGGSGGGSGGMVSK
GEAVIKEFMRFKVHMEGSMNGHEFEIEGEGEGRPYEGTQTAKLKVTKGGPLPFSWDILS
PQFMYGSRAFTKHPADIPDYYKQSFPEGFKWERVMNFEDGGAVTVTQDTSLEDGTLIYK
VKLRGTNFPPDGPVMQKKTMGWEASTERLYPEDGVLKGDIKMALRLKDGGRYLADFKTT
YKAKKPVQMPGAYNVDRKLDITSHNEDYTVVEQYERSEGRHSTGGMDELYKGGGGSG
GGSGILWHEMWHEGLEEASRLYFGERNVKGMFEVLEPLHAMMERGPQTLKETSFNQAY
GRDLMEAQEWCRKYMKSGNVKDLLQAWDLYYHVFRRISK

**Supplementary Note 2|** Double-strand donor DNA sequences for CRISPR knock-in editing. The 40-bp left and right homology arm sequences are underlined. SBP and fluorescent tags are shown in blue and green, respectively.

>>KIF5B (SBP-YFP knock-in at N-terminus)

GAGTGGCTCCCGGCCGCCGGCCCCGGCTGCGAGAAccATGGGACACGTTGTTGAAG
GACTGGCTGGGGAACTTGAACAACTTCGTGCACGACTGGAGCATCACCCACAAGGTG
GTGGAGGTGGCAGCGGAGGCGGTTCCGGAGTGAGCAAGGGCGAGGAGCTGTTCAC
CGGGGTGGTGCCCATCCTGGTCGAGCTGGACGGCGACGTAAACGGCCACAAGTTCA
GCGTGTCCGGCGAGGGCGAGGGCGATGCCACCTACGGCAAGCTGACCCTGAAGTTC
ATCTGCACCACCGGCAAGCTGCCCGTGCCCTGGCCCACCCTCGTGACCACCTTCGG
CTACGGCCTGCAGTGCTTCGCCCGCTACCCCGACCACATGAAGCAGCACGACTTCTT
CAAGTCCGCCATGCCCGAAGGCTACGTCCAGGAGCGCACCATCTTCTTCAAGGACGA
CGGCAACTACAAGACCCGCGCCGAGGTGAAGTTCGAGGGCGACACCCTGGTGAACC
GCATCGAGCTGAAGGGCATCGACTTCAAGGAGGACGGCAACATCCTGGGGCACAAG
CTGGAGTACAACTACAACAGCCACAACGTCTATATCATGGCCGACAAGCAGAAGAAC
GGCATCAAGGTGAACTTCAAGATCCGCCACAACATCGAGGACGGCAGCGTGCAGCT
CGCCGACCACTACCAGCAGAACACCCCCATCGGCGACGGCCCCGTGCTGCTGCCCG
ACAACCACTACCTGAGCTACCAGTCCGCCCTGAGCAAAGACCCCAACGAGAAGCGC
GATCACATGGTCCTGCTGGAGTTCGTGACCGCCGCCGGGATCACTCTCGGCATGGA
CGAGCTGGGCGGAGGATCAGGGGGTGGCTCCGGAGGTGCGGACCTGGCCGAGTGC
AACATCAAAGTGATGTGTCGCT

>>DYNC1H1 (SBP-mNeonGreen knock-in at N-terminus)

CTTCTCATCGCTCCTGGAAGGTCCCGAGCGCGACACCATGGGACACGTTGTTGAAGG
ACTGGCTGGGGAACTTGAACAACTTCGTGCACGACTGGAGCATCACCCACAAGGTGG
TGGAGGTGGCAGCGGAGGCGGTTCCGGAGTGAGCAAGGGCGAGGAGGATAACATG
GCCTCTCTCCCAGCGACACATGAGTTACACATCTTTGGCTCCATCAACGGTGTGGAC
TTTGACATGGTGGGTCAGGGCACCGGCAATCCAAATGATGGTTATGAGGAGTTAAAC
CTGAAGTCCACCAAGGGTGACCTCCAGTTCTCCCCCTGGATTCTGGTCCCTCATATC
GGGTATGGCTTCCATCAGTACCTGCCCTACCCTGACGGGATGTCGCCTTTCCAGGCC
GCCATGGTAGATGGCTCCGGCTACCAAGTCCATCGCACAATGCAGTTTGAAGATGGT

GCCTCCCTTACTGTTAACTACCGCTACACCTACGAGGGAAGCCACATCAAAGGAGAG
GCCCAGGTGAAGGGGACTGGTTTCCCTGCTGACGGTCCTGTGATGACCAACTCGCT
GACCGCTGCGGACTGGTGCAGGTCGAAGAAGACTTACCCCAACGACAAAACCATCAT
CAGTACCTTTAAGTGGAGTTACACCACTGGAAATGGCAAGCGCTACCGGAGCACTGC
GCGGACCACCTACACCTTTGCCAAGCCAATGGCGGCTAACTATCTGAAGAACCAGCC
GATGTACGTGTTCCGTAAGACGGAGCTCAAGCACTCCAAGACCGAGCTCAACTTCAA
GGAGTGGCAAAAGGCCTTTACCGATGTGATGGGCATGGACGAGCTGTACAAGGGCG
GAGGATCAGGGGGTGGCTCCGGAGGTTCGGAGCCCGGGGGCGGCGGCGGCGAGG
ACGGCTCGGCCG

>>ARPC3 (mScarlet-SBP knock-in at C-terminus)

GAGACAGTTCATGAACAAGAGTCTTTCAGGACCTGGACAGGGCGGAGGATCAGGGG
GTGGCTCCGGAGGTGTGAGCAAGGGCGAGGCAGTGATCAAGGAGTTCATGCGGTTC
AAGGTGCACATGGAGGGCTCCATGAACGGCCACGAGTTCGAGATCGAGGGCGAGGG
CGAGGGCCGCCCCTACGAGGGCACCCAGACCGCCAAGCTGAAGGTGACCAAGGGT
GGCCCCCTGCCCTTCTCCTGGGACATCCTGTCCCCTCAGTTCATGTACGGCTCCAGG
GCCTTCACCAAGCACCCCGCCGACATCCCCGACTACTATAAGCAGTCCTTCCCCGAG
GGCTTCAAGTGGGAGCGCGTGATGAACTTCGAGGACGGCGGCGCCGTGACCGTGAC
CCAGGACACCTCCCTGGAGGACGGCACCCTGATCTACAAGGTGAAGCTCCGCGGCA
CCAACTTCCCTCCTGACGGCCCCGTAATGCAGAAGAAGACAATGGGCTGGGAAGCG
TCCACCGAGCGGTTGTACCCCGAGGACGGCGTGCTGAAGGGCGACATTAAGATGGC
CCTGCGCCTGAAGGACGGCGGCCGCTACCTGGCGGACTTCAAGACCACCTACAAGG
CCAAGAAGCCCGTGCAGATGCCCGGCGCCTACAACGTCGACCGCAAGTTGGACATC
ACCTCCCACAACGAGGACTACACCGTGGTGGAACAGTACGAACGCTCCGAGGGCCG
CCACTCCACCGGCGGCATGGACGAGCTGTACAAGGGTGGAGGTGGCAGCGGAGGC
GGTTCCGGAGGACACGTTGTTGAAGGACTGGCTGGGGAACTTGAACAACTTCGTGCA
CGACTGGAGCATCACCCACAAGGTTGAAGGGAGCCCGGGCAGCCACCGTCTCCAGA
GCCCTGGG

**Supplementary Video 1|** Time-lapse images of U2OS cells expressing streptavidin condensates encoded by PB1-mCherry-streptavidin (magenta) and SBP-YFP (green). Cells were treated with biotin at time 0. Scale bar: 10 μm.

**Supplementary Video 2|** Time-lapse images of *KIF5B*$^{SBP/SBP}$ cells expressing streptavidin condensates encoded by PB1-mCherry-streptavidin (magenta). Endogenous SBP-YFP-KIF5B is shown in green. Cells were treated with biotin at time 0. Scale bar: 10 μm.

**Supplementary Video 3|** Time-lapse images of *KIF5B*$^{SBP/SBP}$ cells expressing streptavidin condensates encoded by PB1-streptavidin and HaloTag-RAB6A (magenta). Endogenous SBP-YFP-KIF5B is shown in green. Cells were treated with biotin at time 0. Scale bar: 10 μm.

**Supplementary Video 4|** Time-lapse images of *DYNC1H1*$^{SBP/SBP}$ cells expressing streptavidin condensates encoded by PB1-streptavidin and HaloTag-RAB5A (magenta). Endogenous SBP-mNeonGreen-DYNC1H1 is shown in green. Cells were treated with biotin at time 0. Scale bar: 10 μm.

**Supplementary Video 5|** Time-lapse images of *DYNC1H1*$^{SBP/SBP}$ cells expressing streptavidin condensates encoded by PB1-streptavidin and HaloTag-RAB7A (magenta). Endogenous SBP-mNeonGreen-DYNC1H1 is shown in green. Cells were treated with biotin at time 0. Scale bar: 10 μm.

**Supplementary Video 6|** Time-lapse images of *ARPC3*$^{SBP/SBP}$ cells expressing streptavidin condensates encoded by PB1-streptavidin and Lifeact-mNeonGreen (green). Endogenous ARPC3-mScarlet-SBP is shown in magenta. Cells were treated with biotin at time 0. Scale bar: 10 μm.

**Supplementary Video 7|** Time-lapse images of U2OS cells expressing streptavidin condensates encoded by PB1-FKBP, streptavidin-mCerulean3-FRB (cyan) and SBP-YFP (green). The streptavidin-mCerulean3-FRB is shown in magenta in the merged images. Cells were treated with rapamycin and biotin at time 0 and 45 min, respectively. Scale bar: 10 µm.

**Supplementary Video 8|** Time-lapse images of *DYNC1H1*$^{SBP/SBP}$ cells expressing streptavidin condensates encoded by PB1-FKBP, streptavidin-mScarlet-FRB (cyan) and SBP-YFP (green). Cells were treated with rapamycin and biotin at time 0 and 30 min, respectively. Scale bar: 10 µm.
